# Litter Matters: The Importance of Decomposition Products for Soil Bacterial Diversity and abundance of key groups of the N cycle in Tropical Areas

**DOI:** 10.1101/2023.03.03.530969

**Authors:** Priscila Pereira Diniz, Beatriz Maria Ferrari Borges, Aline Pacobahyba de Oliveira, Maurício Rizzato Coelho, Osnar Obede da Silva Aragão, Thiago Gonçalves Ribeiro, Fernando Igne Rocha, Bruno José Rodrigues Alves, Márcia Reed Rodrigues Coelho, Eustáquio Souza Dias, James R. Cole, Adina Chuang Howe, Siu Mui Tsai, Ederson da Conceição Jesus

## Abstract

This study investigated the contribution of soil organic layers to bacterial diversity evaluations. We used a forest in the eastern Amazon and an adjacent pasture as model systems. Distinct organic and organo-mineral layers were identified in the forest and pasture floors, including the litter, partially and wholly decomposed organic material, and the mineral and rhizospheric soils. DNA was extracted, and 16S rRNA gene sequencing and qPCR were performed to assess bacterial community structure and the abundance of critical groups of the N cycle. We observed a clear vertical gradient in bacterial community composition. Species followed a log-normal distribution, with the highest richness and diversity observed in transitional organic layers of both land uses. Generally, critical groups of the N cycle were more abundant in these transitional layers, especially in the pasture’s fragmented litter and in the forest’s partially decomposed organic material. Considering the organic layers increased diversity estimates significantly, with the highest alpha and gamma bacterial diversity observed on the pasture floor and the highest beta diversity on the forest floor. The results show that organic layers harbor significant bacterial diversity in natural and anthropized systems and suggest that they can be crucial for maintaining the N cycle in these ecosystems, highlighting the need to consider them when studying soil bacterial diversity.

## 1. Introduction

The increasing deforestation for pasture and agriculture in the last decades has posed a threat to the Amazon rainforest, with an average annual expansion rate of 4.6% between 1987 and 2017 (Souza et al., 2020). In response to this scenario, several studies have evaluated the impacts of deforestation and land-use change on soil microbial communities, observing significant changes in their structure, composition, and functionality(Jesus et al., 2009; Mendes et al., 2015; Merloti et al., 2019; Mueller et al., 2014; Navarrete et al., 2011; Paula et al., 2014; Rodrigues et al., 2013).

One of the most notable findings is the strong influence of soil pH and its associated variables on the structure and diversity of bacterial communities (Carvalho et al., 2016; Jesus et al., 2009; Rocha et al., 2021; Rodrigues et al., 2013). Another finding is the lower, unexpected diversity in the forest soil compared to areas deforested for pasture and agriculture (Jesus et al., 2009). These findings have been consistently confirmed by subsequent research (Carvalho et al., 2016; Rodrigues et al., 2013) and were systematized by Petersen et al. (2019). Rodrigues et al. (2013) also observed high alpha diversity associated with low beta diversity in pastures, suggesting a possible biotic homogenization and consequent decline in functional diversity, as later studies have also shown (Kroeger et al., 2018; Paula et al., 2014). However, Carvalho et al. (2016) found higher alpha, beta, and gamma diversities in pastures and agricultural soils. These authors attributed the contrasting results to differences in soil and forest types, which is supported by Rocha et al. (2021), who observed greater differences in diversity in naturally low-fertility soils than in naturally high-fertility soils. Thus, the impact of land-use change on bacterial beta diversity in Amazonian soils remains a question that requires further attention.

The majority of studies mentioned above, with a few exceptions (Moreira et al., 2021; Ritter et al., 2018; Rocha et al., 2021), focuses on microbial communities in the mineral portion of superficial soil horizons. The common approach is to remove the litter and sample immediately underneath, which generally corresponds to the section of soil dominated by mineral material (for example, see Carvalho et al., 2016; Khan et al., 2019; Kroeger et al., 2018; Merloti et al., 2019; Navarrete et al., 2015; Rodrigues et al., 2013). However, this approach, which probably was borrowed from soil fertility assessment studies, ignores the greater complexity of forest soils. Recently studies have shown that this focus can lead to misinterpretation and underestimation of microbial diversity (Rocha et al., 2021).

An essential feature of the forest is the accumulation of a significant amount of litter, whose decomposition products provide a substantial source of nutrients (Herrera et al., 1978; Quesada et al., 2011). The roots are often intertwined with organic matter in various stages of decomposition, forming a root mat on top of the mineral soil. This feature has been reported at least since the 1970s (Herrera et al., 1978; Stark and Jordan, 1978) and has been suggested to be a vital nutrient conservation mechanism in the Amazon forest (Jordan; Herrera, 1981). In this environment, there is typically a layer of recently deposited leaves (L), a layer of fragmented material but still recognizable as litter (F), and an organic horizon (O or H) (Ponge, 2003). These layers constitute different environments that can harbor microbial communities with specific compositions and metabolisms (Baldrian et al., 2012; López-Mondéjar et al., 2015; Prescott and Grayston, 2013).

A great amount of information about the soil microbiota is lost by not taking these layers and their differences into account as they may harbor a significant part of the active forest microbiota. Excluding these layers can lead to an underestimation of the relevant microbial diversity related to nutrient cycling on the forest floor (Rocha et al., 2021). Similar to what was found in studies focusing on the mineral portion of the soil, the structure and diversity of microbial communities in these organic layers change in response to factors such as plant diversity and litter quality (Chapman and Newman, 2009; Pei et al., 2017; Santonja et al., 2018; Zheng et al., 2018). In this respect, litter transformation can be detected by ^13^C isotopic abundance, as it tends to increase as CO2 is released into the atmosphere. In addition, these layers are directly impacted by fire (Santos and Nelson, 2013; Kauffman et al., 1995; Uhl et al., 1988), possibly leading to the loss of an essential component of soil microbial communities. Since the litter and other organic layers are the most affected by these fires, their alteration may have a relevant impact on nutrient cycles due to eliminating key microbial groups.

Therefore, it is necessary to understand how rainforest soil bacterial communities contribute to soil microbial diversity and how they are affected by the land cover change. This study worked under the hypotheses that (1) the organic layers of the forest soil contribute to a greater diversity of prokaryotes in this environment and (2) that these layers favor a greater diversity of microorganisms in the cycle of N.

## 2. Material and Methods

### 2.1 Characterization of the study area and sampling

The study was carried out in the District of São Joaquim do Ituquara, municipality of Baião, in the state of Pará, Brazil (Supplementary Figure 1). Samples were taken from a primary forest and an adjacent pasture with *Urochola* sp. (*syn. Brachiaria* sp.) in December 2017. A detailed description of the sites can be found in the Supplementary Material.

A transect of 250 m was delimited in each land use system, with five sampling points equally spaced (50 m) (Supplementary Figure 1). The distance between the closest points of the forest and the pasture was 100 m. Composite samples were collected at each point with a 25 cm x 25 cm metallic square (Figure 1). Each composite sample consisted of three subsamples, collected either in exclusively organic sections or dominated by organic material (hereinafter called organic layers), as well as in the sections where there is a predominance of mineral material (hereinafter referred to as mineral soil).

**Figure 1.**
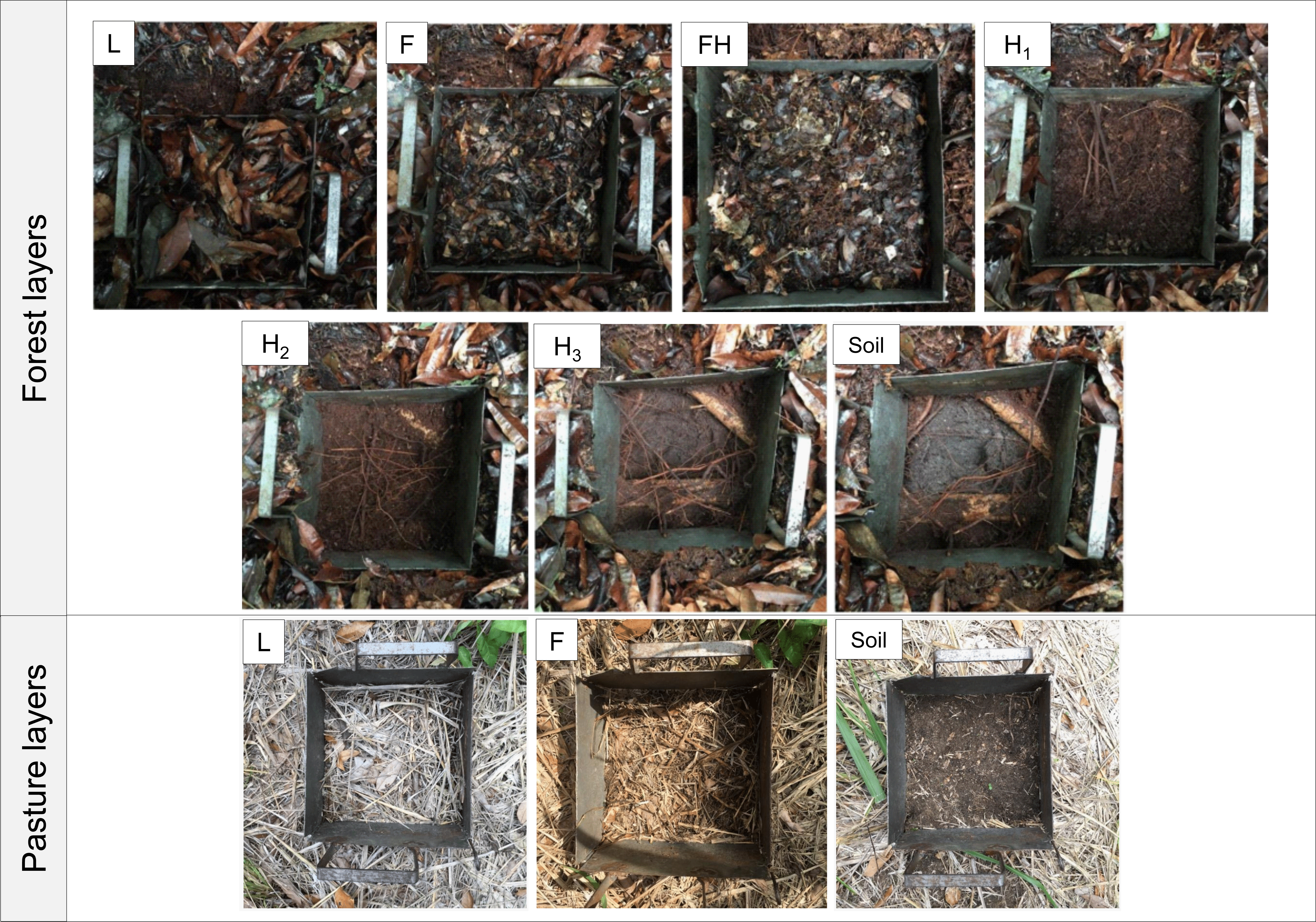
Sampling of organic layers and mineral soil from the forest and pasture with the aid of a metallic template.

The organic layers, which correspond to the O (organic) horizon, were defined as L, F, and H layers (Santos et al., 2018) and arbitrarily subdivided into sublayers FH; H_1_, H_2_, and H_3_ based on their decomposition stage and morphological differentiation. The sampling procedure was extended for another 10 cm below the organic layers to characterize the mineral portion of the soil (Figure 1), which corresponds to part of the A horizon. The way of distinguishing and collecting the material was based on the classification of humus forms and the visual examination of the sampled material [22, 35]. In total, seven distinct layers were sampled in the forest (L, F, FH, H_1_, H_2_, H_3_, and the mineral soil), and four in the pasture (L, F, and the mineral soil), including the rhizosphere soil, corresponding to the soil adhering to the roots of grasses (*Urochloa* sp.).

After sampling, the samples were sieved in a 2 mm mesh (except L and F) and, later, stored at -80°C. The transition layer FH was subdivided into FH, containing sieved material < 2mm, and FHc, containing coarse material retained in the sieve. All samples were used for chemical characterization as described in the Supplementary Material.

### 2.4 DNA Extraction

L, F, and FH samples containing leaf and root fragments were macerated in liquid nitrogen and 0.25 g of the macerated material was used for DNA extraction with the DNeasy Power Soil kit (Qiagen, Hilden, Germany) according to the manufacturer’s recommendations. The samples from the other layers were used directly for DNA extraction with the same kit, and 0.50 g of sample material was used for extractions.

The DNA obtained was quantified in a Nanodrop 2000c spectrophotometer (Thermo Fisher Scientific Inc., USA), observing its purity by the A260/280 ratio and its integrity by electrophoresis in a 1% agarose gel (100 V for 30 min), stained with ethidium, with visualization of the DNA bands in a transilluminator.

### 2.5 Bioinformatics sequencing and analysis

Large-scale sequencing of the bacterial 16S rRNA gene was performed by the Institute for Genomics and Systems Biology Next Generation Sequencing (IGSB-NGS) at the Argonne National Laboratory in Chicago, USA, following the amplification protocols of the Earth Microbiome Project (Thompson et al., 2017). The primer pair 515F and 806R (Caporaso et al., 2011), which amplifies the V4 region of the 16S rRNA, was used. Sequencing was performed using the Illumina MiSeq platform (2 × 250 bp paired-end). The raw sequence data is available in the SRA database of Genbank with the accession number PRJNA927008.

The sequences obtained were analyzed using the DADA2 pipeline (1.8) (Callahan et al., 2016) incorporated into the R software. The sequences were demultiplexed, filtered for quality (Q score greater than 20, with a minimum length of 250 bp in the forward sequences and 225 bp in the reverse sequences), joined, and classified using the SILVA v.138 database (Quast et al., 2013). Finally, the generated ASV table was converted into a phyloseq object using the phyloseq package (McMurdie and Holmes, 2013) for further analysis.

### 2.6 Real-time PCR of nitrogen cycle genes

The abundances of Bacteria, Archaea, and nitrogen cycle functional genes were determined by quantitative real-time PCR (qPCR) on gene copy number per gram of substrate (soil or organic material). The following genes were evaluated: 16S rRNA from Bacteria and Archaea; nitrogenase (*nifH*) for nitrogen fixation; ammonium monooxygenase (*amoA*) from Bacteria and Archaea for nitrification (NH_3_ to NH_2_OH); nitrite reductase (*nirS*; NO_2_^-^ to NO) and nitrous oxide reductase (*nosZ*; N_2_O to N_2_) for denitrification. Analyzes were performed using the StepOnePlus™ Real-Time PCR system with 96-well plates (Applied Biosystems, Foster City, CA, USA). From serial dilutions, standard curves of genes of microorganisms known to contain N cycle enzymes, previously amplified by PCR, were constructed. All genes used in the curves were purified with the IllustraTM GFXTM PCR DNA and Gel Band Purification Kit (GE Healthcare, UK) for better efficiency of the standard curves, according to the manufacturer’s recommendations. The microorganisms used as standard, primers, and amplification conditions are shown in Supplementary Table 1. All samples were quantified in triplicate in a final volume of 10 μL of reaction, composed of 5.0 μL of SYBR Green ROX qPCR Master Mix (Thermo Scientific), 0.67 to 1.0 μL of each primer (see Supplementary Table 1), 0.5 μL of bovine serum albumin (BSA, 6 mg.mL^- 1^), 1.0 μL of DNA diluted 50 times and enough water for PCR to complete the final volume. For all assays, amplification efficiency ranged from 96% to 106%, and R^2^ values were 0.98–0.99.

### 2.7 Data analysis and statistics

The analyzes were performed using the R software using packages phyloseq, vegan, easyanova, and agricolae (Maintainer and Arnhold, 2022; McMurdie and Holmes, 2013; Mendiburu, 2021; Oksanen et al., 2005). A filtering procedure was applied to remove low-prevalence sequences and sequences from eukaryotes, chloroplasts, and mitochondria. The length of the species distribution gradient was verified by detrended correspondence analysis (DCA). Non-metric multidimensional scaling (NMDS) was used to visualize dissimilarities in bacterial community structure based on the Bray-Curtis distance over the abundance of ASVs per sample.

Permutational multivariate analysis of variance (PERMANOVA) (Anderson, 2001) was applied to observe possible significant differences in the structure of bacterial communities, also based on the Bray-Curtis distance matrix. To assess the community phylum composition, the relative abundance data were checked for normality (Shapiro- Wilk) and homogeneity of variance (Bartlett). The relative abundance data followed the assumption of non-parametric tests, and the Kruskal-Wallis test was performed at 5% significance with the FDR (False Discovery Rate) adjustment method.

A sample containing 7,007 readings was removed from the analysis, and the data were rarefied for the smallest number of reads (10,402 reads). Alpha diversity measures (Chao1 and Shannon) were calculated separately for each of the organic layers and the mineral soil and for all layers belonging to the same land use system. The significance was tested with the non-parametric Kruskal-Wallis test at 5% significance with the FDR (False Discovery Rate) adjustment method.

Diversity partition analysis was performed using Hill numbers through the Entropart package as explained elsewhere (Chao et al., 2014; Daly et al., 2018; Jost, 2006; Marcon and Hérault, 2015; Rocha et al., 2021). Diversity metrics were calculated for all layers separately and after associating all layers in a single system. Preemption, log-Normal, Zipf, and Zipf-Mandelbrot species abundance models were tested using the radfit command of the vegan package (Oksanen et al., 2005). The comparison of these models was based on the Akaike Information Criterion (AIC), a measure of the relative quality of a statistical model.

The abundances of the functional genes of the N cycle, obtained through qPCR, were analyzed in the program StepOne Software v2.3 (Applied Biosystem, USA) for their specificity and efficiency. Data were compared using the Kruskal-Wallis test at 5% significance with the FDR correction method for multiple comparisons.

## 3. Results

### 3.1 Chemical properties of soils

Soil C:N ratios decreased with depth in both land uses, more abruptly from L to H_1_ on the forest floor and from L to F on the pasture floor (Supplementary Figure 2).

The δ^13^C and δ ^15^N values on the forest floor became more positive with depth (r = 0.962; p>0.001). The same pattern was observed on the pasture floor, where the bulk and rhizosphere soils showed the same C:N ratio and δ^15^N, while a difference existed for values of δ^13^C. For the forest floor, the K, Ca, Mg, Cu, Mn, and Zn concentrations also decreased in the direction of more transformed material, from the L layer to mineral soil, while the Fe concentration increased. P concentration was the highest in the litter (L, F, and FH) and the lowest in the organic horizon (H_1_ and H_2_). In the pasture, the Ca and Mg concentrations were higher in L when compared to the F layer (Supplementary Table 2).

The H_3_ layer in the forest and the mineral and rhizospheric soils in the pasture (Supplementary Figure 3) are acidic, but the pH was significantly lower in the forest soil, with an average value of 4.3 against 5.7 and 5.8 in the pasture. The pasture soil showed higher Ca, Fe, Mn, and Zn concentrations, as opposed to the forest floor, which in turn showed higher concentrations of Al and organic C.

### 3.2 Microbial community structure and composition

Forest and pasture bacterial communities were grouped into two distinct groups in the NMDS analysis (stress = 0.0811; Figure 2). A gradient of distribution and substitution of species from the uppermost L layer towards the mineral soil was observed in both land uses. This gradient was long in both systems, with 6.08 and 4.68 standard deviations in forest and pasture, respectively, clearly showing the existence of an ecocline and a high beta-diversity (Supplementary Figure 3). The species abundance distribution model with the best fit, both the forest and pasture floors, was the log- normal, followed by the Mandelbrot and the Zipf (Figure 3).

**Figure 2.**
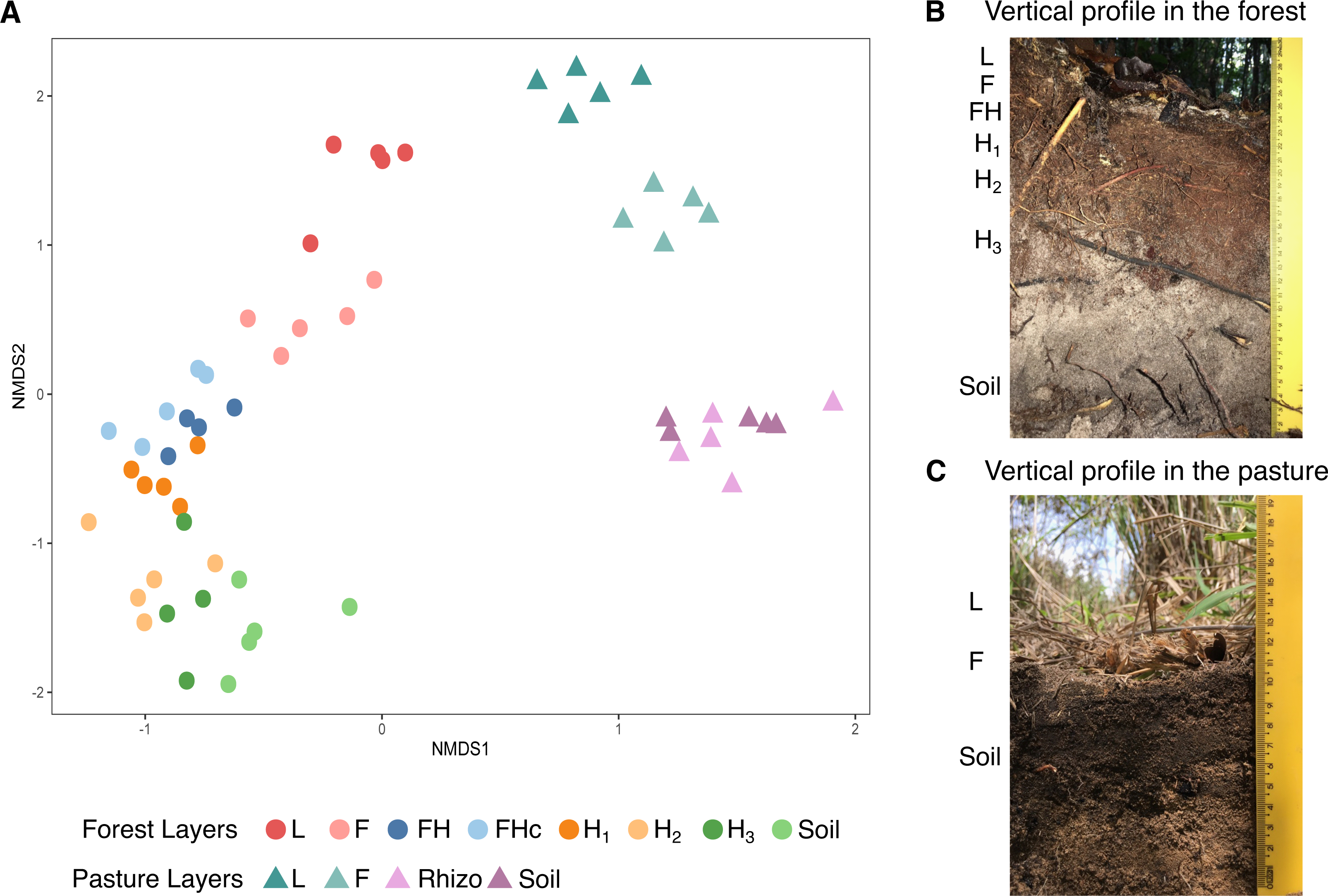
Non-metric multidimensional scaling analysis (NMDS) of bacterial communities from the organic and organomineral layers of forest and pasture (A); soil profile photo under forest (B) and pasture (C).

**Figure 3.**
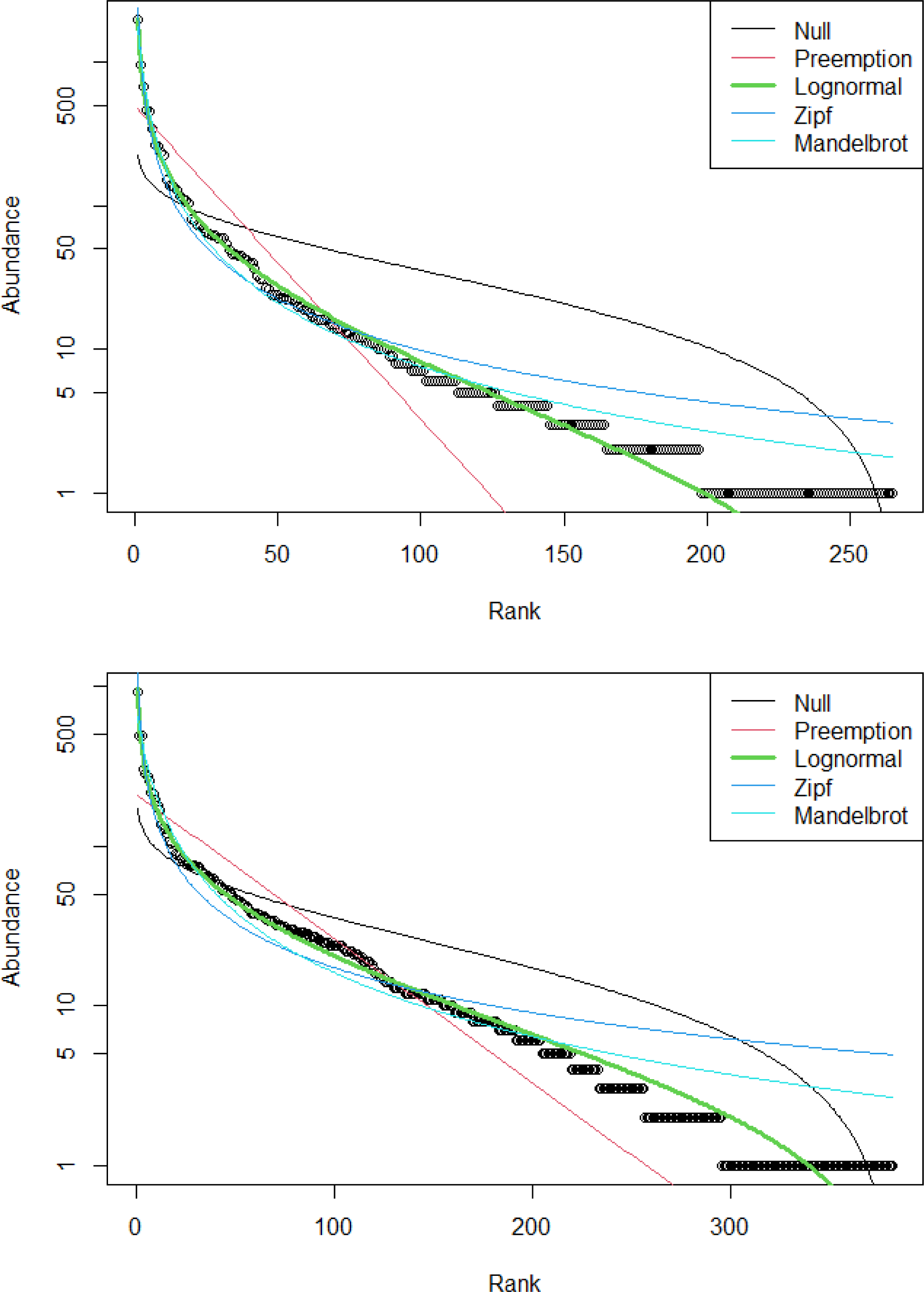
Best-fit of abundance species models to bacterial communities in the forest (top) and pasture (bottom) floors.

PERMANOVA (F = 6.43, P < 0.001) allowed the identification of six different community structures in the forest: (1) L, (2) F, (3) FH/FHc, (4) FH/H1, (5) H_2_/H_3_, and (6) H_3_/mineral soil. Three distinct groups were identified in the pasture: (1) L, (2) F, and (3) mineral and rhizospheric soils. It is worth highlighting that the L layers were those with the smallest dissimilarity between the two land use systems.

Some gradients were observed in the relative abundance of major phyla. The abundances of *Proteobacteria*, *Bacteroidota*, and *Firmicutes* decreased with depth in the forest floor. The last two phyla were not observed from the H_2_ and FH layers down, respectively, if considered an abundance greater than 1% (Figure 4A; Supplementary Figure 4). *Acidobacteriota, Actinobacteriota, Planctomycetota*, and *Verrucomicrobiota* showed increasing abundances with depth and were especially abundant in the organic horizon. In the pasture, the gradients were not as clear due to the smaller number of layers. *Proteobacteria*, *Actinobacteriota*, and *Bacteroidota* were more abundant in the litter (L and F) (Figure 4B). On the other hand, *Firmicutes, Verrucomicrobiota*, and *Planctomycetota* were more abundant in mineral and rhizospheric soils. No difference in abundance was observed for *Acidobacteriota*.

**Figure 4.**
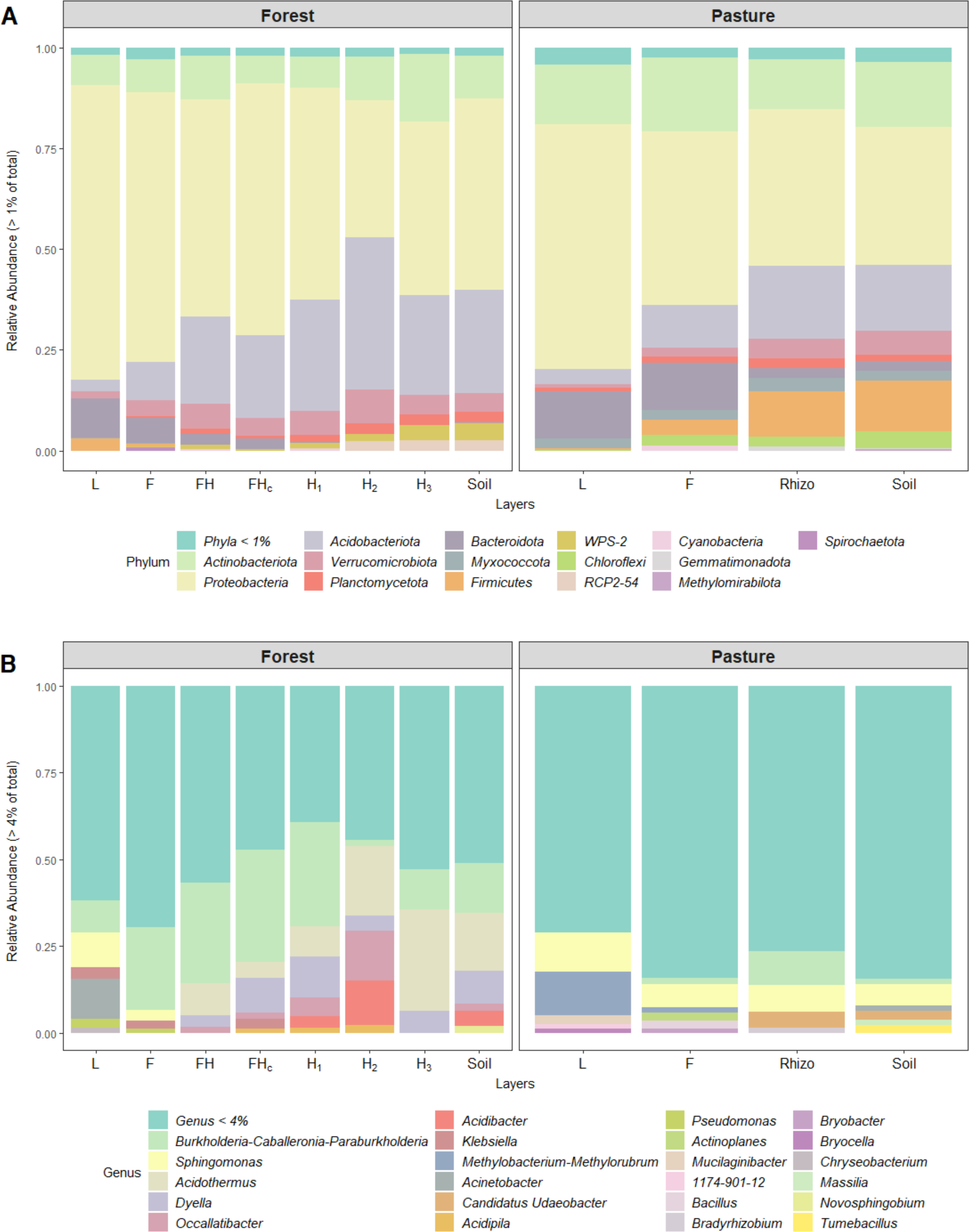
Relative abundance of bacterial phyla (A) and (B) genera in organic layers at different stages of decomposition and mineral soil of forest and pasture. L entire leaves; F, fragmented leaves; FH, fine fragments of the mixture of fragmented leaves and humus; FHc, fragments greater than 2 mm from the mixture of fragmented leaves and humus; H_1_, humus with fine roots; H_2_, humus; H_3_, humus mixed with soil mineral material; Rhizo, rhizospheric soil; Soil, initial 10 cm of mineral soil from forest or pasture.

Forty-five percent of the forest sequences and 82% of those in the pasture account for genera with an abundance of less than 4%, indicating the great microbial diversity at this level, especially in the pasture (Figure 4B). Overall, *Burkholderia- Caballeronia-Paraburkholderia* were the most abundant genera in the forest, accounting for 20% of the sequences. *Sphingomonas* (9%) was the most abundant genus in the pasture. In the forest, *Acinetobacter* predominates in the L layer (11%), *Burkholderia-Caballeronia-Paraburkholderia* in F, FH, FHc, and H_1_ (24%; 29%, 32%; and 29%, respectively) and *Acidothermus* in H_2_, H_3_, and the mineral soil (20%, 30%, and 20%, respectively). In the pasture, *Methylobacterium-Methylorubrum* prevailed in L (3%), *Sphingomonas* in F and the mineral soil (7% in each), and *Burkholderia- Caballeronia-Paraburkholderia* in the rhizospheric soil (10%).

### 3.4 Diversity of microbial community

The bacterial richness in the forest differed significantly between the H_2_ and FHc only (H= 14.66, p= 0.041; Figure 5A). L showed the lowest richness in the pasture, while the other layers did not differ from each other (H = 8.17, p = 0.042) (Figure 5B). When considering the communities of all layers together, the overall richness of the pasture floor was higher than that of the forest floor (H = 29.90, p = 4.5e-08) (Figure 5C).

**Figure 5.**
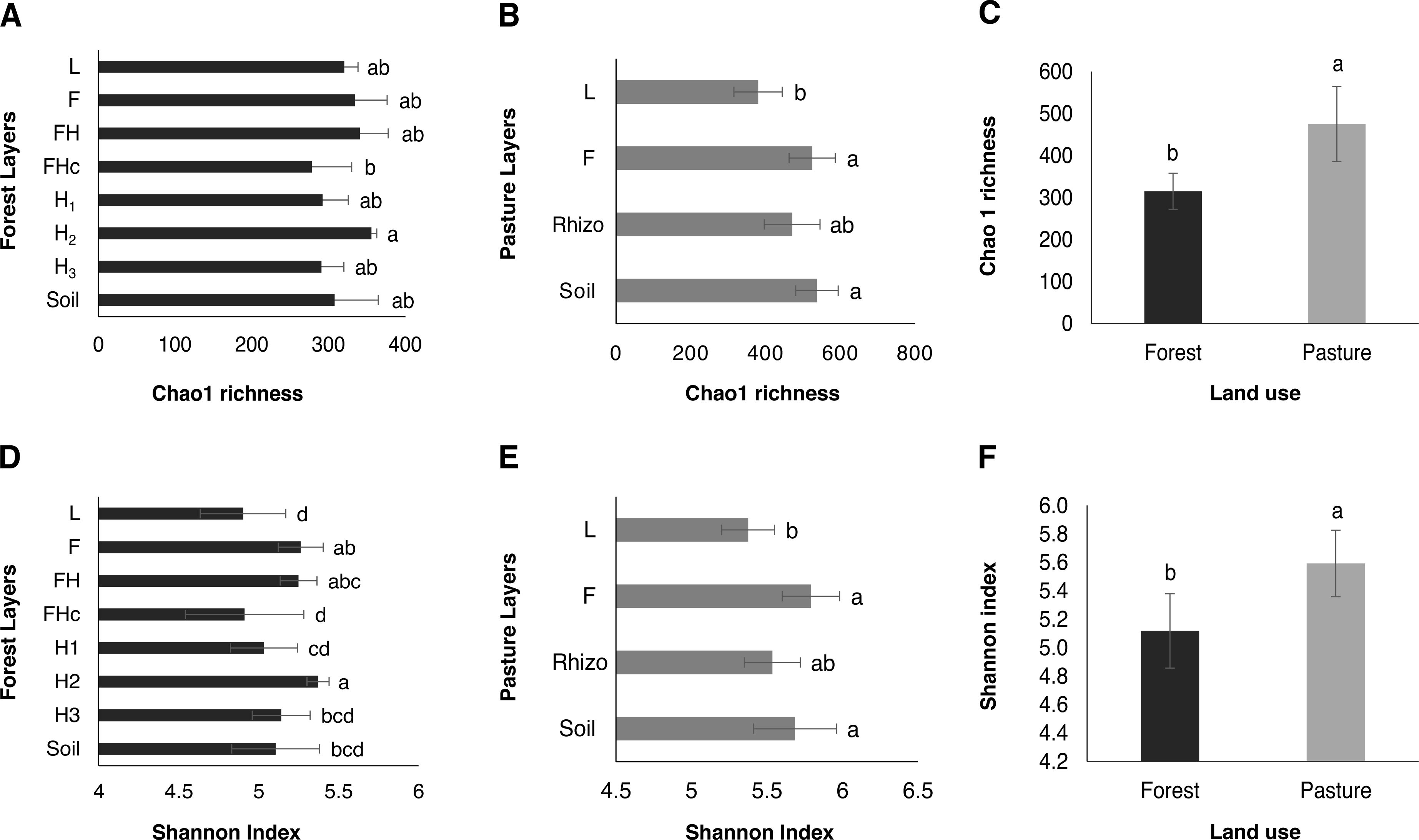
Chao1 richness and Shannon index of bacterial communities in organic layers in different stages of decomposition and mineral soil (A and D, respectively) of forest, (B, E) pasture and (C, F) total land use systems. Error bars indicate standard deviation of five independent replicates. Different letters refer to the statistical difference between organic layers and mineral soil or land use system based on the Kruskal Wallis test (P < 0.05). L, entire leaves; F, fragmented leaves; FH, fine fragments of the mixture of fragmented leaves and humus; FHc, fragments greater than 2 mm from the mixture of fragmented leaves and humus; H_1_, humus with fine roots; H_2_, humus; H_3_, humus mixed with soil mineral material; Rhizo, rhizospheric soil; Soil, initial 10 cm of mineral soil from forest or pasture.

In the forest, the Shannon index was the highest in H_2_, the only layer to differ from H_3_ and the mineral soil (H = 19.91, p = 0.0058) (Figure 5D). The L layer showed the highest diversity in the pasture but did not differ from mineral and rhizospheric soils (H = 9.69; p = 0.0214) (Figure 5E). As observed for richness, the Shannon diversity was higher on the pasture floor than on the forest floor (H = 28.25, p = 1.1e-07) (Figure 5F).

The diversity partition analysis showed that for the alpha, beta, and gamma diversities of the forest, the ASVs richness (q = 0) and Shannon exponential (q = 1) were higher for the forest floor in relation to the individual layers (Figure 6A). On the pasture floor, alpha and gamma diversities were also superior to those in the individual layers alone (Figure 6B), but the same was not observed for the beta-diversity.

**Figure 6.**
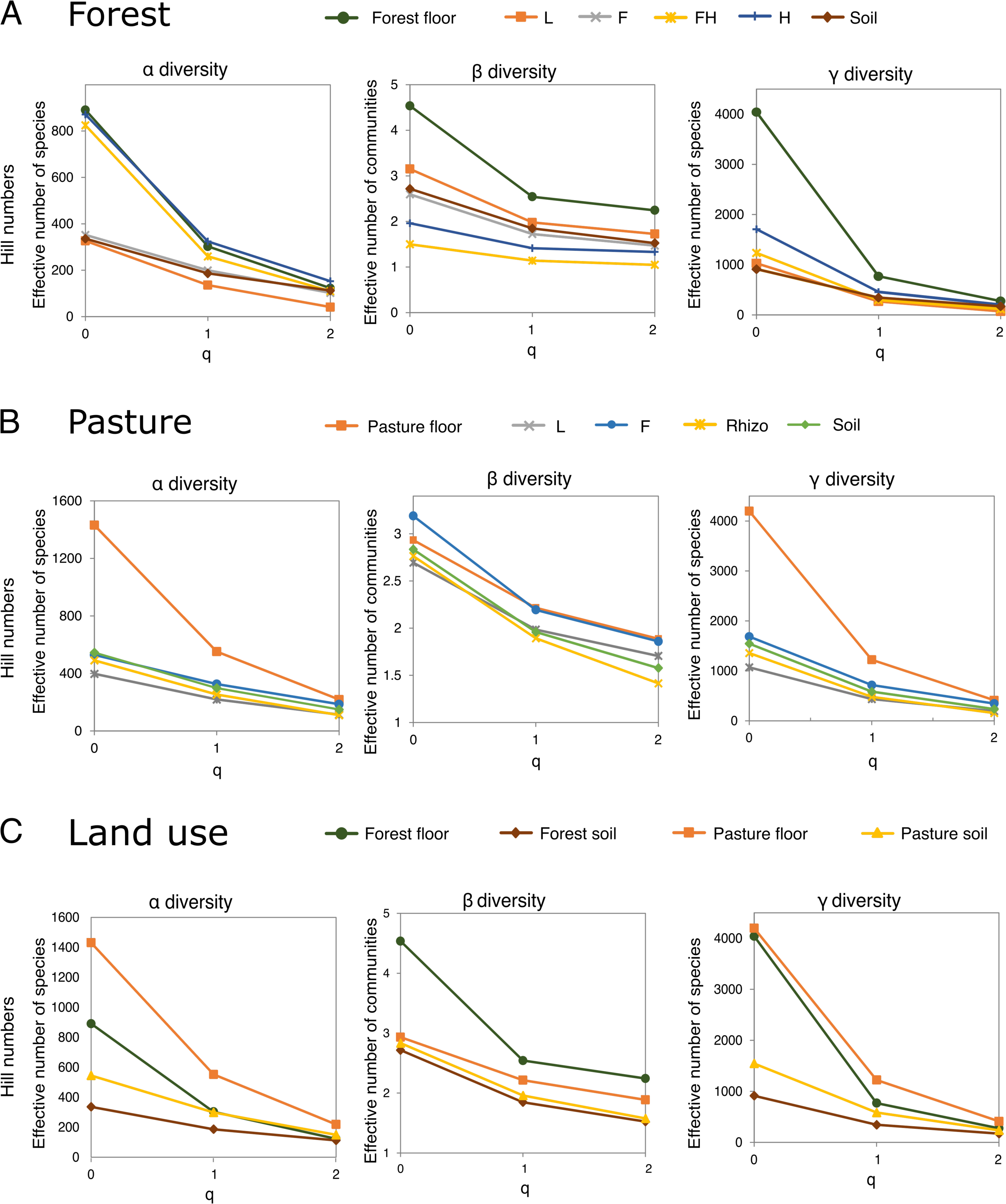
Diversity partition analysis evaluating alpha, beta, and gamma diversities for bacterial communities in forest and pasture, in which q = 0 represents the ASV richness; q = 1, the exponential of Shannon entropy for equally weighted ASVs; q = 2, the inverse of Simpson’s index for dominant rates. The forest floor pasture floors represent the analysis of all layers of the use system together (organic and organominerals); L, entire leaves; F, fragmented leaves; FH, fine and coarse fragments of the mixture of fragmented leaves and humus; H, junction of layers H_1_ (humus with fine roots), H_2_ (humus) and H_3_ (humus mixed with soil mineral material); Rhizo, rhizospheric soil; Soil, initial 10 cm of mineral soil from forest or pasture.

Subsequently, when comparing the two land use systems (Figure 6C), it was observed that in the alpha and gamma diversity scale, the pasture floor was highest than the forest floor and forest mineral soil for the parameters of ASV richness (q = 0) and Shannon exponential (q = 1). However, the beta diversity was higher in the forest floor compared to the pasture floor in all diversity components (Figure 6C). Simpson dominance (q = 2), representing the effective number of dominant species or communities, was similar across all study layers at the alpha and gamma diversity scales. However, on the beta diversity scale, higher values of dominance were recorded for the microbial communities of the forest and pasture floors, with the forest floor being significantly higher than the pasture floor.

### 3.5 Abundance of functional nitrogen cycle genes

In the forest, the abundances of Bacteria were higher in organic layers, especially in FH and H_2_ (H = 36.65, p = 5.4e-06) (Figure 7). In the pasture, layer F held the highest abundance of Bacteria (H = 16.35, p = 0.001) at a level comparable to that of layers FH and H_2_ of the forest. Archaea, in general, were more abundant in the forest compared to the pasture (Figure 7). Like Bacteria, this domain was more abundant in the organic layers, especially FH and H_2_ of the forest (H = 31.74, p = 4.5e-05) and in the F layer of the pasture (H = 16.35, p = 0.001). It is worth noting that the abundances of both domains decreased in the H_1_ layer and increased in the H_2_ layer. In addition, the abundances of organisms from both domains were significantly lower in the mineral soil of both systems (Figure 7).

**Figure 7.**
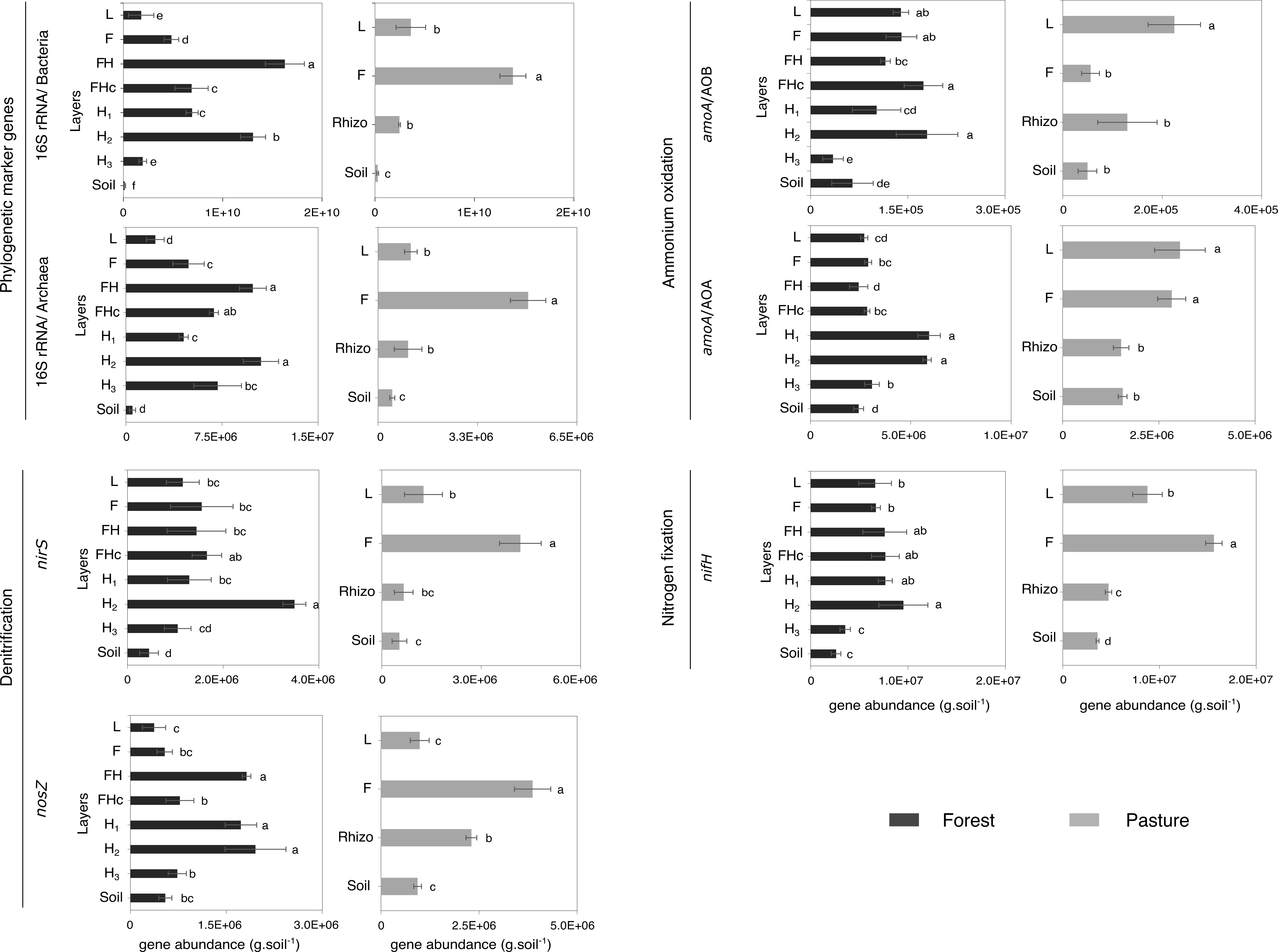
Abundance of Bacteria and Archaea (16S rRNA gene), N_2_-fixing bacteria (*nifH*), ammonia-oxidizing bacteria and archaea (*amoA*), and denitrifying bacteria (*nirS* and *nosZ*) in organic layers at different stages of decomposition and in the mineral soil of the forest and pasture. Error bars indicate standard deviation of five independent replicates. Different letters refer to the statistical difference between organic layers and mineral soil of the same land use system based on the Kruskal Wallis test (P < 0.05). L entire leaves; F, fragmented leaves; FH, fine fragments of the mixture of fragmented leaves and humus; FHc, fragments greater than 2 mm from the mixture of fragmented leaves and humus; H_1_, humus with fine roots; H_2_, humus; H_3_, humus mixed with soil mineral material; Rhizo, rhizospheric soil; Soil, initial 10 cm of mineral soil from forest or pasture.

In the forest, the abundance of the *nifH* gene was higher in the organic layers, especially in the H_2_ layer (H = 26.33, p = 0.00044) (Figure 7). This gene was significantly less abundant in the transitional H_3_ layer and the mineral soil. The organic layers of the pasture also showed a higher *nifH* abundance (H = 17.91, p = 0.00046), mainly on the F layer. Its abundance was lower in the L layer, rhizospheric soil, and the mineral soil (Figure 7).

Ammonia-oxidizing archaea (AOA) were more abundant in the forest H_1_ and H_2_ layers and less in the mineral soil and the L and F layers (H = 29.77, p = 0.00010) (Figure 7). Ammonia-oxidizing bacteria (AOB) were present in considerable abundance in the L and F layers, in addition to being prominent in FHc and H_2_ (H = 29.38, p = 0.00012) (Figure 7). These groups were also abundant in the pasture L and F organic layers. AOA was also abundant in the L and F layers (H = 14.46, p = 0.0023), while AOB stood out in the L layer only (H = 12.60, p = 0.0056).

The *nirS* denitrification gene was more abundant in the H_2_ layer of the forest, making its presence quite different from the other layers (H = 25.05, p = 0.0007) (Figure 7). In the pasture, *nirS* stood out in the F layer, which was significantly superior to L and mineral and rhizospheric soils (H = 13.67, p = 0.0034). The *nosZ* gene was more abundant in the layers FH, H_1,_ and H_2_ of the forest, and its lowest values were associated with mineral soil and organic layers L and F (H = 28.85, p = 0.0001) (Figure 7). In the pasture, *nosZ* stood out in the F layer and was less abundant in the mineral soil and the L layer (H = 16.26, p = 0.0010).

## 4. Discussion

Little attention has been paid to the microbiota of organic materials commonly found on the soil surface under forests and pastures in the tropics. Therefore, changes in tropical soil microbial diversity due to land use and deforestation have been studied with a focus on soil mineral horizons, disregarding the litter and its transformation products, with rare exceptions (Ritter et al., 2018; Rocha et al., 2021). Here, an alternative and holistic approach to address this topic is proposed. A similar approach has been used to study microbial diversity in temperate forests (Trap et al., 2011) and soil fauna focusing on humus forms (Izadi et al., 2017; Salmon, 2018; Salmon et al., 2005), which represent the set of organic and organo-mineral layers in soil (Embrapa, 1997; Ponge, 2003).

In this work, we observe a clear gradient of microbial community structure along the forest profile, following the stages of organic matter decomposition. Additionally, the log-normal distribution was the underlying species distribution, implying the existence of a hierarchical niche structure with several independent environmental factors acting on species abundances (May, 1975; Sugihara, 1980). The bacterial species are substituted along the vertical profile, indicating the decomposition nature of the organic material as a critical factor defining community composition, with the different layers constituting different micro-habitats and creating an ecocline, as shown by the long species distribution gradients. High microbial diversity was observed in the H layer, especially in what we defined as layer H_2_. This layer consists largely of wet organic material with little or no visible plant structure (except fine roots) because of the action of the edaphic fauna and fungi, which transform the litter and make nutrients available to plants and microorganisms. The high presence of fine roots in this layer has already been reported as a mechanism of nutrient conservation in the Amazon Forest in low-fertility soils (Herrera et al., 1978). As the litter is decomposed, the quality of the substrate is modified, changing the parameters that lead to the differentiation of microbial communities such as C:N and C:P ratios, moisture content, pH, and proportions of cellulose, hemicellulose, and lignin (Brockett et al., 2012; Papa et al., 2014; Uhl et al., 1988). Thus, these parameters act as ecological filters, likely resulting in the functional compartmentalization of microbial communities. The higher δ^13^C in layers with highly decomposed organic matter and in the mineral soil suggests that the recalcitrant material, such as lignin, seems not to stabilize in soil, as lignin is generally depleted in δ^13^C but will influence microbial diversity (Wedin et al., 1995). Microbial diversity also seems related to the enriched δ^13^C in rhizosphere soil compared to bulk mineral soil in the pasture area. It is likely related to a more significant input of C_4_-C through decaying roots and exudates. Nonetheless, the resulting 30% more replacement of forest carbon by pasture carbon can be attributable to shifts in microorganism composition.

A gradient in community composition has also been observed in the pasture floor, but with well-defined boundaries between litter and soil communities. Bacterial diversity was especially high in the F layer and, surprisingly, equal to those of the rhizospheric soil, a region considered a hotspot of microbial diversity and activity (Hinsinger et al., 2009; Kuzyakov and Blagodatskaya, 2015; Steinauer et al., 2016). However, these two compartments differed considerably as shown by their clear difference in microbial community composition. This layer was noticeably wetter compared to the underlying L layer, a factor that might have favored microbial diversity and abundance (Brockett et al., 2012), and its decomposition also makes a large amount of organic C available (Papa et al., 2014). There was considerable similarity between L communities from both land uses, reinforcing the interpretation of an influence by the nature of the substrate. Although the plant species compositions and diversity levels are distinct, the nature of the material (plant vs. soil material) contributed to selecting closely related species.

We tested if the pattern of higher alpha diversity of soil bacteria under pasture and agriculture (Carvalho et al., 2016; Jesus et al., 2009; Navarrete et al., 2015; Pedrinho et al., 2019; Rodrigues et al., 2013) would remain if the organic layers were considered. The alpha diversity of the pasture floor was significantly higher than that of the forest floor, supporting the pattern of higher soil bacterial diversity in pastures.

However, beta diversity was higher in the forest floor if compared to the pasture’s mineral soil as previously observed by Rocha et al. (2021). This highlights the increment in diversity caused by adding the organic layers in these analyses. In fact, comparing the diversity values of the mineral soil from both sites with those in the forest floor and pasture floor, a considerable increment was observed at all diversity scales for both land uses, except for the pasture’s beta diversity. Thus, intrinsic diversity in the organic layers is underestimated with the current collection methodology. No significant difference was observed for beta diversity when comparing the mineral soils from both areas. There is still no consensus on how land use change may affect soil beta diversity and whether changes may be characteristic of the study site. Increases (Carvalho et al., 2016; Rocha et al., 2021) and decreases (Rodrigues et al., 2013) in beta diversity have already been observed in pastures in different regions of the Amazon.

Parameters such as pH and plant diversity have already been suggested as possible predictors of beta diversity (Carvalho et al., 2016; Prober et al., 2015). When taking the whole forest and pasture floors into account, no difference in beta diversity was observed, although a higher effective number of communities was recorded in the forest, especially when considering the ASV richness (q = 0). The increment in beta diversity caused by the addition of the organic layers in the forest may indicate plant diversity’s influence on the heterogeneity of soil microbial communities (Mitchell et al., 2010; Prober et al., 2015; Zinger et al., 2011). The gamma diversity was similar on both floors and higher than the values found for their respective mineral soils. This similarity in beta and gamma diversity values between the land use systems, although not indicating a biotic homogenization as in Rodrigues et al. (2013) does not exclude the possibility of reduction of essential microbial functions and ecosystem resilience because, as previously discussed, the microbial community structure of the forest is different from that observed in the pasture.

The enrichment in ^15^N with the aging of organic material as observed here is a natural process related to N transformation and losses (Högberg, 1997), indicating that differences between forest and pasture floors should be relevant for N transformations. For this reason, we focused on N-cycle microorganisms as a critical group for nutrient cycling in these environments. Bacteria, Archaea, and key groups in the N cycle were especially abundant in the organic layers in both land use systems. In the forest, the H_2_ layer is associated with the highest absolute abundances of all functional genes, as is the pasture’s F layer (except for AOB, which is more abundant in L). These results indicate that the significant N transformations may occur more intensively above the mineral soil. It is recognized that gene quantification does not reflect the total biological potential for N cycling. Still, as observed for microbial diversity, these layers stand out as potential key compartments and therefore deserve full attention in microbial ecology studies.

The litter mineralization, plant-microorganism interactions, and the interaction among the key groups of the N-cycle may have been important factors for the enrichment of functional genes in these organic layers. The high abundance of diazotrophs, AOA, and AOB in the organic layers may also be related to the presence of these groups in the phyllosphere of the forest canopy and pasture plants, where the occurrence of free-living nitrogen fixation and nitrification has also been detected (Fürnkranz et al., 2008; Guerrieri et al., 2020, 2015; Moreira et al., 2021; Papen et al., 2002; Stanton et al., 2019; Van Langenhove et al., 2020; Watanabe et al., 2016). In this way, N transformations in the plant canopy also represent a critical unquantified above- ground compartment and act as a constant microbial inoculum for the litter. In the Amazon, free-living nitrogen fixation has already been observed as higher in litter than in mineral soil (Moreira et al., 2021), although these rates are still controversial for other environments(Reed et al., 2008; Van Langenhove et al., 2021, 2020). Regarding nitrification, studies that relate this process to litter are rare (Adams, 1986). However, it is known that some grasses can emit N_2_O from their leaves, probably through the transport of gas from the soil (Bowatte et al., 2014). In turn, AOB have already been observed emitting N_2_O in pasture leaves (Bowatte et al., 2014). Given this, if AOA and AOB abundances are higher in organic layers, greenhouse gas emission rates may be underestimated by excluding litter in both agricultural and pasture systems.

Low O_2_ concentration, readily available organic compounds as an energy source and the presence of nitrate are factors that favor denitrification (Galland et al., 2019). These factors can be found in the rhizosphere (Achouak et al., 2019). However, interestingly, in the pasture, *nirS* and *nosZ* were more abundant in the litter, which could also influence greenhouse gas emissions. The highest *nirS/nosZ* ratios (Supplementary Figure 8) were related to the layers composed of leaves in both environments, indicating that the litter may be responsible for higher N_2_O emissions via denitrification compared to the other soil layers. N_2_O emissions have already been correlated with *nirS* abundance (Morales et al., 2010; Rasche et al., 2010).

## 5. Conclusion

Soil organic layers both in the forest and pasture harbor considerable bacterial diversity, and their exclusion from soil biodiversity studies can underestimate the diversity of relevant bacterial groups contributing for nutrient cycling in natural and anthropized environments. These layers also harbor abundant populations of microorganisms of the N cycle, reinforcing their potential contribution for nutrient cycling in the soil. We suggest that more attention should be paid to these organic layers in soil microbial ecology studies since they are undoubtedly fundamental for maintaining these ecosystems.

## Acknowledgments

We acknowledge the USAID and the National Academies of Sciences, Engineering, and Medicine of the United States (NAS) for funding our research under PEER project 4299, USAID agreement AID-OAA-A-11-00012. Any opinions, findings, conclusions, or recommendations expressed here are those of the authors alone and do not necessarily reflect the views of USAID or the NAS. We also thank the Coordination of Superior Level Improvement – Capes (finance code 001) for the research fellowship provided to Priscila Pereira Diniz, and CNPq for the research fellowships provided to Ederson da Conceição Jesus (project 312781/2022-9) and Fernando Igne Rocha (165571/2017-9). Finally, we thank Mr. João Balieiro de Souza Aragão and Mrs. Maria Mirtes Lopes da Silva, who provided us access to the sampling sites and kindly supported us with our sampling.

## Data Availability Statement

The raw sequence data is available in the SRA database of Genbank with the accession number PRJNA927008. The raw metadata used for statistical analysis are available from the corresponding author on reasonable request.

## References

Achouak, W., Abrouk, D., Guyonnet, J., Barakat, M., Ortet, P., Simon, L., Lerondelle, C., Heulin, T., El Zahar Haichar, F., 2019. Plant hosts control microbial denitrification activity. FEMS Microbiol. Ecol. 95, 21. https://doi.org/10.1093/FEMSEC/FIZ021

Adams, J.A., 1986. Nitrification and ammonification in acid forest litter and humus as affected by peptone and ammonium-n amendment. Soil Biol. Biochem. 18, 45–51. https://doi.org/10.1016/0038-0717(86)90102-1

Anderson, M.J., 2001. A new method for non-parametric multivariate analysis of variance. Austral Ecol. 26, 32–46. https://doi.org/10.1111/J.1442-9993.2001.01070.PP.X

Baldrian, P., Kolaiřík, M., Štursová, M., Kopecký, J., Valášková, V., Větrovský, T., Žifčáková, L., Šnajdr, J., Rídl, J., Vlček, Č., Voříšková, J., 2012. Active and total microbial communities in forest soil are largely different and highly stratified during decomposition. ISME J. 6, 248–258. https://doi.org/10.1038/ismej.2011.95

Bowatte, S., Newton, P.C.D., Brock, S., Theobald, P., Luo, D., 2014. Bacteria on leaves: a previously unrecognised source of N2O in grazed pastures. ISME J. 2015 91 9, 265–267. https://doi.org/10.1038/ismej.2014.118

Brockett, B.F.T., Prescott, C.E., Grayston, S.J., 2012. Soil moisture is the major factor influencing microbial community structure and enzyme activities across seven biogeoclimatic zones in western Canada. Soil Biol. Biochem. 44, 9–20. https://doi.org/10.1016/J.SOILBIO.2011.09.003

Callahan, B.J., McMurdie, P.J., Rosen, M.J., Han, A.W., Johnson, A.J.A., Holmes, S.P., 2016. DADA2: High-resolution sample inference from Illumina amplicon data. Nat. Methods 13, 581–583. https://doi.org/10.1038/nmeth.3869

Caporaso, J.G., Lauber, C.L., Walters, W.A., Berg-Lyons, D., Lozupone, C.A., Turnbaugh, P.J., Fierer, N., Knight, R., 2011. Global patterns of 16S rRNA diversity at a depth of millions of sequences per sample. Proc. Natl. Acad. Sci. U. S. A. 108, 4516–4522.

Carvalho, T.S. de, Jesus, E. da C., Barlow, J., Gardner, T.A., Soares, I.C., Tiedje, J.M., Moreira, F.M. de S., 2016. Land use intensification in the humid tropics increased both alpha and beta diversity of soil bacteria. Ecology 97, 2760–2771. https://doi.org/10.1002/ECY.1513

Chao, A., Chiu, C.H., Jost, L., 2014. Unifying Species Diversity, Phylogenetic Diversity, Functional Diversity, and Related Similarity and Differentiation Measures Through Hill Numbers. https://doi.org/10.1146/annurev-ecolsys-120213-091540 45, 297–324. https://doi.org/10.1146/ANNUREV-ECOLSYS-120213-091540

Chapman, S.K., Newman, G.S., 2009. Biodiversity at the plant–soil interface: microbial abundance and community structure respond to litter mixing. Oecologia 162, 763– 769. https://doi.org/10.1007/S00442-009-1498-3

Daly, A.J., Baetens, J.M., De Baets, B., 2018. Ecological Diversity: Measuring the Unmeasurable. Math, 6, 119. https://doi.org/10.3390/MATH6070119

Empresa Brasileira de Pesquisa Agropecuária – EMBRAPA. 1997. Manual de métodos de análise de solo. 2sd ed. Centro Nacional de Pesquisas de Solos, Rio de Janeiro.

Fürnkranz, M., Wanek, W., Richter, A., Abell, G., Rasche, F., Sessitsch, A., 2008. Nitrogen fixation by phyllosphere bacteria associated with higher plants and their colonizing epiphytes of a tropical lowland rainforest of Costa Rica. ISME J. 2008 25 2, 561–570. https://doi.org/10.1038/ismej.2008.14

Galland, W., Piola, F., Burlet, A., Mathieu, C., Nardy, M., Poussineau, S., Blazère, L., Gervaix, J., Puijalon, S., Simon, L., Haichar, F. el Z., 2019. Biological denitrification inhibition (BDI) in the field: A strategy to improve plant nutrition and growth. Soil Biol. Biochem. 136, 107513. https://doi.org/10.1016/J.SOILBIO.2019.06.009

Guerrieri, R., Lecha, L., Mattana, S., Cáliz, J., Casamayor, E.O., Barceló, A., Michalski, G., Peñuelas, J., Avila, A., Mencuccini, M., 2020. Partitioning between atmospheric deposition and canopy microbial nitrification into throughfall nitrate fluxes in a Mediterranean forest. J. Ecol. 108, 626–640. https://doi.org/10.1111/1365-2745.13288

Guerrieri, R., Vanguelova, E.I., Michalski, G., Heaton, T.H.E., Mencuccini, M., 2015. Isotopic evidence for the occurrence of biological nitrification and nitrogen deposition processing in forest canopies. Glob. Chang. Biol. 21, 4613–4626. https://doi.org/10.1111/GCB.13018

Herrera, R., Klinge, H., Medina, E., 1978. Amazon ecosystems: their structure and functioning with particular emphasis on nutrients. Interciencia 3, 223–231.

Hinsinger, P., Bengough, A.G., Vetterlein, D., Young, I.M., 2009. Rhizosphere: biophysics, biogeochemistry and ecological relevance. Plant Soil 321, 117–152. https://doi.org/10.1007/S11104-008-9885-9

Högberg, P., 1997. Tansley Review No. 95 15N natural abundance in soil–plant systems. New Phytol. 137, 179–203. https://doi.org/10.1046/J.1469-8137.1997.00808.X

Izadi, M., Habashi, H., Waez-Mousavi, S.M., 2017. Variation in soil macro-fauna diversity in seven humus orders of a Parrotio-Carpinetum forest association on Chromic Cambisols of Shast-klateh area in Iran. Eurasian Soil Sci. 50, 341–349. https://doi.org/10.1134/S106422931703005X

Jesus, E.D.C., Marsh, T.L., Tiedje, J.M., Moreira, F.M.D.S., 2009. Changes in land use alter the structure of bacterial communities in Western Amazon soils. ISME J 3, 1004–1011. https://doi.org/10.1038/ismej.2009.47

Jordan, C.F., Herrera, R., 1981. Tropical Rain Forests: Are Nutrients Really Critical? The America n Naturalist. 17:7–13. https://doi.org/10.1086/283696

Jost, L., 2006. Entropy and diversity. Oikos 113, 363–375. https://doi.org/10.1111/J.2006.0030-1299.14714.X

Kauffman, J.B., Cummings, D.L., Ward, D.E., Babbitt, R., 1995. Fire in the Brazilian Amazon: 1. Biomass, nutrient pools, and losses in slashed primary forests. Oecologia 104, 397–408. https://doi.org/10.1007/BF00341336

Khan, M.A.W., Bohannan, B.J.M., Nüsslein, K., Tiedje, J.M., Tringe, S.G., Parlade, E., Barberán, A., Rodrigues, J.L.M., 2019. Deforestation impacts network co- occurrence patterns of microbial communities in Amazon soils. FEMS Microbiol. Ecol. 95. https://doi.org/10.1093/FEMSEC/FIY230

Kroeger, M.E., Delmont, T.O., Eren, A.M., Meyer, K.M., Guo, J., Khan, K., Rodrigues, J.L.M., Bohannan, B.J.M., Tringe, S.G., Borges, C.D., Tiedje, J.M., Tsai, S.M., Nüsslein, K., 2018. New biological insights into how deforestation in amazonia affects soil microbial communities using metagenomics and metagenome- assembled genomes. Front. Microbiol. 9, 1635. https://doi.org/10.3389/fmicb.2018.01635

Kuzyakov, Y., Blagodatskaya, E., 2015. Microbial hotspots and hot moments in soil: Concept & review. Soil Biol. Biochem. 83, 184–199. https://doi.org/10.1016/J.SOILBIO.2015.01.025

López-Mondéjar, R., Voříšková, J., Větrovský, T., Baldrian, P., 2015. The bacterial community inhabiting temperate deciduous forests is vertically stratified and undergoes seasonal dynamics. Soil Biol. Biochem. 87, 43–50. https://doi.org/10.1016/J.SOILBIO.2015.04.008

Maintainer, E.A., Arnhold, E., 2022. Package “easyanova” Title Analysis of Variance and Other Important Complementary Analyses.

Marcon, E., Hérault, B., 2015. entropart: An R Package to Measure and Partition Diversity. J. Stat. Softw. 67, 1–26. https://doi.org/10.18637/JSS.V067.I08

May, R.M., 1975. Patterns of species abundance and diversity, in: University, H. (Ed.), Ecology and Evolution of Communities. Cambridge 81–120.

McMurdie, P.J., Holmes, S., 2013. Phyloseq: An R Package for Reproducible Interactive Analysis and Graphics of Microbiome Census Data. PLoS One 8, e61217. https://doi.org/10.1371/journal.pone.0061217

Mendes, L.W., Tsai, S.M., Navarrete, A.A., de Hollander, M., van Veen, J.A., Kuramae, E.E., 2015. Soil-Borne Microbiome: Linking Diversity to Function. Microb. Ecol. 70, 255–265. https://doi.org/10.1007/S00248-014-0559-2

Mendiburu, F. 2021. Package ‘agricolae’ version 1.3–5.

Merloti, L.F., Mendes, L.W., Pedrinho, A., de Souza, L.F., Ferrari, B.M., Tsai, S.M., 2019. Forest-to-agriculture conversion in Amazon drives soil microbial communities and N-cycle. Soil Biol. Biochem. 137, 107567. https://doi.org/10.1016/J.SOILBIO.2019.107567

Mitchell, R.J., Hester, A.J., Campbell, C.D., Chapman, S.J., Cameron, C.M., Hewison, R.L., Potts, J.M., 2010. Is vegetation composition or soil chemistry the best predictor of the soil microbial community? Plant Soil 333, 417–430. https://doi.org/10.1007/S11104-010-0357-7

Morales, S.E., Cosart, T., Holben, W.E., 2010. Bacterial gene abundances as indicators of greenhouse gas emission in soils. ISME J. 4, 799–808. https://doi.org/10.1038/ismej.2010.8

Moreira, J.C.F., Brum, M., de Almeida, L.C., Barrera-Berdugo, S., de Souza, A.A., de Camargo, P.B., Oliveira, R.S., Alves, L.F., Rosado, B.H.P., Lambais, M.R., 2021. Asymbiotic nitrogen fixation in the phyllosphere of the Amazon forest: Changing nitrogen cycle paradigms. Sci. Total Environ. 773, 145066. https://doi.org/10.1016/J.SCITOTENV.2021.145066

Mueller, R.C., Paula, F.S., Mirza, B.S., Rodrigues, J.L., Nüsslein, K., Bohannan, B.J., 2014. Links between plant and fungal communities across a deforestation chronosequence in the Amazon rainforest. ISME J. 8, 1548–1550. https://doi.org/10.1038/ismej.2013.253

Navarrete, A.A., Taketani, R.G., Mendes, L.W., Cannavan, F.S., Moreira, F.M.S., Tsai, S.M., 2011. Land-use systems affect archaeal community structure and functional diversity in western amazon soils. Rev. Bras. Cienc. do Solo 35, 1527–1540. https://doi.org/10.1590/s0100-06832011000500007

Navarrete, A.A., Tsai, S.M., Mendes, L.W., Faust, K., De Hollander, M., Cassman, N.A., Raes, J., Van Veen, J.A., Kuramae, E.E., 2015. Soil microbiome responses to the short-term effects of Amazonian deforestation. Mol. Ecol. 24, 2433–2448. https://doi.org/10.1111/MEC.13172

Oksanen, J., Kindt, R., O’, B., Maintainer, H., 2005. The vegan Package Title Community Ecology Package.

Papa, S., Cembrola, E., Pellegrino, A., Fuggi, A., Fioretto, A., 2014. Microbial enzyme activities, fungal biomass and quality of the litter and upper soil layer in a beech forest of south Italy. Eur. J. Soil Sci. 65, 274–285. https://doi.org/10.1111/EJSS.12112

Papen, H., Geßler, A., Zumbusch, E., Rennenberg, H., 2002. Chemolithoautotrophic Nitrifiers in the Phyllosphere of a Spruce Ecosystem Receiving High Atmospheric Nitrogen Input. Curr. Microbiol. 44, 56–60. https://doi.org/10.1007/S00284-001-0074-9

Paula, F.S., Rodrigues, J.L.M., Zhou, J., Wu, L., Mueller, R.C., Mirza, B.S., Bohannan, B.J.M., Nüsslein, K., Deng, Y., Tiedje, J.M., Pellizari, V.H., 2014. Land use change alters functional gene diversity, composition and abundance in Amazon forest soil microbial communities. Mol. Ecol. 23, 2988–2999. https://doi.org/10.1111/MEC.12786

Pedrinho, A., Mendes, L.W., Merloti, L.F., Fonseca, M.C., Cannavan, F.S., Tsai, S.M., 2019. Forest-to-pasture conversion and recovery based on assessment of microbial communities in Eastern Amazon rainforest. FEMS Microbiol. Ecol. 95, 236. https://doi.org/10.1093/FEMSEC/FIY236

Pei, Z., Leppert, K.N., Eichenberg, D., Bruelheide, H., Niklaus, P.A., Buscot, F., Gutknecht, J.L.M., 2017. Leaf litter diversity alters microbial activity, microbial abundances, and nutrient cycling in a subtropical forest ecosystem. Biogeochem. 134, 163–181. https://doi.org/10.1007/S10533-017-0353-6

Petersen, I.A.B., Meyer, K.M., Bohannan, B.J.M., 2019. Meta-Analysis Reveals Consistent Bacterial Responses to Land Use Change Across the Tropics. Front. Ecol. Evol. 7, 391. https://doi.org/10.3389/FEVO.2019.00391/BIBTEX

Ponge, J.F., 2003. Humus forms in terrestrial ecosystems: a framework to biodiversity. Soil Biol. Biochem. 35, 935–945. https://doi.org/10.1016/S0038-0717(03)00149-4

Prescott, C.E., Grayston, S.J., 2013. Tree species influence on microbial communities in litter and soil: Current knowledge and research needs. For. Ecol. Manage. 309, 19– 27. https://doi.org/10.1016/j.foreco.2013.02.034

Prober, S.M., Leff, J.W., Bates, S.T., Borer, E.T., Firn, J., Harpole, W.S., Lind, E.M., Seabloom, E.W., Adler, P.B., Bakker, J.D., Cleland, E.E., Decrappeo, N.M., Delorenze, E., Hagenah, N., Hautier, Y., Hofmockel, K.S., Kirkman, K.P., Knops, J.M.H., La Pierre, K.J., Macdougall, A.S., Mcculley, R.L., Mitchell, C.E., Risch, A.C., Schuetz, M., Stevens, C.J., Williams, R.J., Fierer, N., 2015. Plant diversity predicts beta but not alpha diversity of soil microbes across grasslands worldwide. Ecol. Lett. 18, 85–95. https://doi.org/10.1111/ELE.12381

Quast, C., Pruesse, E., Yilmaz, P., Gerken, J., Schweer, T., Yarza, P., Peplies, J., Glöckner, F.O., 2013. The SILVA ribosomal RNA gene database project: improved data processing and web-based tools. Nucleic Acids Res. 41, D590– D596. https://doi.org/10.1093/NAR/GKS1219

Quesada, C.A., Lloyd, J., Anderson, L.O., Fyllas, N.M., Schwarz, M., Czimczik, C.I., 2011. Soils of Amazonia with particular reference to the RAINFOR sites. Biogeosciences 8, 1415–1440. https://doi.org/10.5194/BG-8-1415-2011

Rasche, F., Knapp, D., Kaiser, C., Koranda, M., Kitzler, B., Zechmeister-Boltenstern, S., Richter, A., Sessitsch, A., 2010. Seasonality and resource availability control bacterial and archaeal communities in soils of a temperate beech forest. ISME J. 2011 53 5, 389–402. https://doi.org/10.1038/ismej.2010.138

Reed, S.C., Cleveland, C.C., Townsend, A.R., 2008. TREE SPECIES CONTROL RATES OF FREE-LIVING NITROGEN FIXATION IN A TROPICAL RAIN FOREST. Ecology 89, 2924–2934. https://doi.org/10.1890/07-1430.1

Ritter, C.D., Zizka, A., Roger, F., Tuomisto, H., Barnes, C., Nilsson, R.H., Antonelli, A., 2018. High-throughput metabarcoding reveals the effect of physicochemical soil properties on soil and litter biodiversity and community turnover across Amazonia. PeerJ 2018, e5661. https://doi.org/10.7717/PEERJ.5661/SUPP-10

Rocha, F.I., Ribeiro, T.G., Fontes, M.A., Schwab, S., Coelho, M.R.R., Lumbreras, J.F., da Motta, P.E.F., Teixeira, W.G., Cole, J., Borsanelli, A.C., Dutra, I. dos S., Howe, A., de Oliveira, A.P., Jesus, E. da C., 2021c. Land-Use System and Forest Floor Explain Prokaryotic Metacommunity Structuring and Spatial Turnover in Amazonian Forest-to-Pasture Conversion Areas. Front. Microbiol. 12, 909. https://doi.org/10.3389/FMICB.2021.657508/BIBTEX

Rodrigues, J.L.M., Pellizari, V.H., Mueller, R., Baek, K., Jesus, E.C., Paula, F.S., Mirza, B., Hamaou, G.S., Tsai, S.M., Feiglf, B., Tiedje, J.M., Bohannan, B.J.M., Nusslein, K., 2013a. Conversion of the Amazon rainforest to agriculture results in biotic homogenization of soil bacterial communities. PNAS 110, 988–993. https://doi.org/10.1073/pnas.1220608110

Salmon, S., 2018. Changes in humus forms, soil invertebrate communities and soil functioning with forest dynamics. Appl. Soil Ecol. 123, 345–354. https://doi.org/10.1016/J.APSOIL.2017.04.010

Salmon, S., Geoffroy, J.J., Ponge, J.F., 2005. Earthworms and collembola relationships: effects of predatory centipedes and humus forms. Soil Biol. Biochem. 37, 487– 495. https://doi.org/10.1016/J.SOILBIO.2004.08.011

Santonja, M., Foucault, Q., Rancon, A., Gauquelin, T., Fernandez, C., Baldy, V., Mirleau, P., 2018. Contrasting responses of bacterial and fungal communities to plant litter diversity in a Mediterranean oak forest. Soil Biol. Biochem. 125, 27–36. https://doi.org/10.1016/J.SOILBIO.2018.06.020

Santos, A.R., Nelson, B.W., 2013. Leaf decomposition and fine fuels in floodplain forests of the Rio Negro in the Brazilian Amazon. J. Trop. Ecol. 29, 455–458. https://doi.org/10.1017/S0266467413000485

Santos, R.D., Jacomine, P.T.K., Anjos, L.H.C., Oliveira, V.A., Lumbreras, J.F., Coelho, M.R., Cunha, T.J.F., Oliveira, J.B., 2018. Sistema Brasileiro de Classificação de Solos, 5th ed. Brasília, DF.

Souza, C.M., Shimbo, J.Z., Rosa, M.R., Parente, L.L., Alencar, A.A., Rudorff, B.F.T., Hasenack, H., Matsumoto, M., Ferreira, L.G., Souza-Filho, P.W.M., de Oliveira, S.W., Rocha, W.F., Fonseca, A. V., Marques, C.B., Diniz, C.G., Costa, D., Monteiro, D., Rosa, E.R., Vélez-Martin, E., Weber, E.J., Lenti, F.E.B., Paternost, F.F., Pareyn, F.G.C., Siqueira, J. V., Viera, J.L., Neto, L.C.F., Saraiva, M.M., Sales, M.H., Salgado, M.P.G., Vasconcelos, R., Galano, S., Mesquita, V. V., Azevedo, T., 2020. Reconstructing three decades of land use and land cover changes in brazilian biomes with landsat archive and earth engine. Remote Sens. 12, 2735. https://doi.org/10.3390/RS12172735

Stanton, D.E., Batterman, S.A., Von Fischer, J.C., Hedin, L.O., 2019. Rapid nitrogen fixation by canopy microbiome in tropical forest determined by both phosphorus and molybdenum. Ecology 100, e02795. https://doi.org/10.1002/ECY.2795

Stark, N.M., Jordan, C.F., 1978. Nutrient Retention by the Root Mat of an Amazonian Rain Forest. Ecology 59, 434–437. https://doi.org/10.2307/1936571

Steinauer, K., Chatzinotas, A., Eisenhauer, N., 2016. Root exudate cocktails: the link between plant diversity and soil microorganisms? Ecol. Evol. 6, 7387–7396. https://doi.org/10.1002/ECE3.2454

Sugihara, G., 1980. Minimal Community Structure: An Explanation of Species Abundance Patterns. https://doi.org/10.1086/283669 116, 770–787. https://doi.org/10.1086/283669

Thompson, L.R., Sanders, J.G., McDonald, D. et al. 2017. A communal catalogue reveals Earth’s multiscale microbial diversity. Nature 551, 457–463. https://doi.org/10.1038/nature24621

Trap, J., Laval, K., Akpa-Vinceslas, M., Gangneux, C., Bureau, F., Decaëns, T., Aubert, M., 2011. Humus macro-morphology and soil microbial community changes along a 130-yr-old Fagus sylvatica chronosequence. Soil Biol. Biochem. 43, 1553–1562. https://doi.org/10.1016/J.SOILBIO.2011.04.005

Uhl, C., Kauffman, J.B., Cummings, D.L., 1988. Fire in the Venezuelan Amazon 2: Environmental Conditions Necessary for Forest Fires in the Evergreen Rainforest of Venezuela. Oikos 53, 176. https://doi.org/10.2307/3566060

Van Langenhove, L., Depaepe, T., Verryckt, L.T., Fuchslueger, L., Donald, J., Leroy, C., Krishna Moorthy, S.M., Gargallo-Garriga, A., Ellwood, M.D.F., Verbeeck, H., Van Der Straeten, D., Peñuelas, J., Janssens, I.A., 2021. Comparable canopy and soil free-living nitrogen fixation rates in a lowland tropical forest. Sci. Total Environ. 754, 142202. https://doi.org/10.1016/J.SCITOTENV.2020.142202

Van Langenhove, L., Depaepe, T., Vicca, S., van den Berge, J., Stahl, C., Courtois, E., Weedon, J., Urbina, I., Grau, O., Asensio, D., Peñuelas, J., Boeckx, P., Richter, A., Van Der Straeten, D., Janssens, I.A., 2020. Regulation of nitrogen fixation from free-living organisms in soil and leaf litter of two tropical forests of the Guiana shield. Plant Soil 450, 93–110.

Watanabe, K., Kohzu, A., Suda, W., Yamamura, S., Takamatsu, T., Takenaka, A., Koshikawa, M.K., Hayashi, S., Watanabe, M., 2016. Microbial nitrification in throughfall of a Japanese cedar associated with archaea from the tree canopy. Springerplus 5, 1–15.

Wedin, D.A., Tieszen, L.L., Dewey, B., Pastor, J., 1995. Carbon Isotope Dynamics During Grass Decomposition and Soil Organic Matter Formation. Ecology 76, 1383–1392. https://doi.org/10.2307/1938142

Zheng, H., Chen, Y., Liu, Y., Zhang, J., Yang, W., Yang, L., Li, H., Wang, L., Wu, F., Guo, L., 2018. Litter quality drives the differentiation of microbial communities in the litter horizon across an alpine treeline ecotone in the eastern Tibetan Plateau. Sci. Reports 8, 1–11. https://doi.org/10.1038/s41598-018-28150-1

Zinger, L., Lejon, D.P.H., Baptist, F., Bouasria, A., Aubert, S., Geremia, R.A., Choler, P., 2011. Contrasting Diversity Patterns of Crenarchaeal, Bacterial and Fungal Soil Communities in an Alpine Landscape. PLoS One 6, e19950. https://doi.org/10.1371/JOURNAL.PONE.0019950

