## Supplementary material for "Litter Matters: The Importance of Decomposition Products for Soil Bacterial Diversity and abundance of key groups of the N cycle in Tropical Areas": Suplemental files

### Supplementary methods

#### Characterization of the study area and soils

The site is in an extensive alluvial plain of predominantly flat relief, which transitions to gently waved relief in the vicinity of the thalwegs. The sampling points under the forest are close and practically perpendicular to the stream, with sandy soils classified as typical Orthic Humiluvic Spodosols according to the Brazilian Soil Classification System (SiBCS) (Santos et al., 2018).The pasture is in a higher position in the landscape and is located in typical Dystrophic Yellow Latosols of medium texture [1]. Two trenches were opened in each land use system to classify the soils. The soils were sampled and classified according to Santos et al. (Santos et al., 2018) and Santos et al. (Santos et al., 2015). Soil drilling using an auger was carried out along the transects to verify the soil attributes' homogeneity for their classification. The area is homogeneous, keeping the same soil class in each land use system studied. The region's climate is classified as tropical monsoon (Am – Köppen classification), with an average temperature of 26°C and an average annual rainfall of 2,202 mm.

#### Brief description of the sampled material

The organic layers, which correspond to the O (organic) horizon, were defined as L, F, and H layers according to the SiBCS (Santos et al., 2018) and subdivided into sublayers FH, $H_1$, $H_2$, and $H_3$. These subdivisions are not defined in the cited manual; they were arbitrarily defined based on their decomposition stage and morphological differentiation. Layer L consists of freshly deposited and non-fragmented litter leaves. The F consists of fragmented leaves and small amounts of fine organic material. FH is characterized by an abundance of roots mixed with fragmented leaves and a greater amount of fragmented organic matter with a diameter smaller than 2 mm. Layer H was subdivided into three sublayers according to their morphological differentiation: the first, $H_1$, is characterized by the abundance of fine fragmented organic material, with a high abundance of tree roots and other forest plants; the second, $H_2$, has visibly a lower abundance of roots; $H_3$, constitutes a transition between the organic layer and the mineral soil (organomineral layer), with a mixture containing higher organic matter values. Finally, the mineral soil, of organomineral constitution, but with a predominance of the properties of mineral constituents (mineral material, according to Santos et al., 2018), was characterized by a primarily sandy texture in both study areas. This section corresponds to the most superficial part of the surface horizon, of moderate type A, according to SiBCS (Santos et al., 2018), with average organic carbon content, grayish or darker (black), and thickness generally equal to or greater than 20 cm.

**Chemical analysis of organic layers and mineral soil**

The chemical analyzes were carried out following the recommendations of Embrapa (Embrapa, 1997). Organic samples (L, F, FH, FHc, $H_{1,}$ and $H_2$ from the forest and L and F from the pasture) were initially submitted to digestion in a heating block (except for potassium analysis, which was carried out by microwave digestion). Nitrogen was quantified by the Kjeldahl method with steam distillation and potassium by flame photometer. The other elements (P, Ca, Mg, Cu, Fe, Mn, and Zn) were analyzed by atomic absorption spectrometry.

The following analyses were carried out with samples from $H_3$ and mineral and rhizospheric soils: organic carbon determination by the potassium dichromate method in sulfuric medium and titration; and total nitrogen by the Kjeldahl method. The micronutrients Cu, Fe, Mn, and Zn were extracted by Mehlich-1 and determined by atomic absorption spectrometry. The $Ca^{2+}$, $Mg^{2+,}$ and $Al^{3+}$ were extracted using a 1 mol $L^{-1}$ KCl solution. For H + Al, a 0.5 mol $L^{-1}$ calcium acetate solution at pH 7.0 was used. Na+, K+, and P extractions were carried out with a solution of 0.0125 mol $L^{-1}$ + HCl 0.05 mol $L^{-1}$. $Ca^{2+}$ and $Mg^{2+}$ contents were determined by atomic absorption spectroscopy. $K^+$ and $Na^+$ contents were determined by flame photometry. P content was determined by colorimetry, and $Al^{3+}$ and H+Al by titration. The sum of exchangeable bases (ES) was calculated as the sum of $Ca^{2+}$, $Mg^{2+}$, $K^+$, and $Na^+$. The cation exchange capacity (CEC) was calculated by the sum of ES and potential acidity (H + Al). Base saturation (V%) was calculated as the ratio between exchangeable cations and CEC multiplied by 100.

Analysis of the natural abundance of isotopes $^{13}C$ and $^{15}N$ was performed for all samples by the John Day Laboratory of Embrapa Agrobiologia (Seropédica, RJ, Brazil). The samples were dried at 60ºC, homogenized, and ground to a fine powder using a ball mill. Afterward, the samples were weighed in tin capsules in triplicate and analyzed in a mass ratio spectrometer for stable isotopes (Finnigan MAT, Bremen, Germany).

**Supplementary Table 1** – Primers, standard strains, and amplification conditions of N-cycle genes for qPCR analysis.

| Target genes | Standard | Primer | Sequence (5' - 3') | Fragment length (pb) | Primer volume (µL) | Primer concentration (pmol) | Primer reference | Amplification condition |
|---|---|---|---|---|---|---|---|---|
| *16S rRNA* Bacteria | DSM 50090 *Pseudomonas fluorescens* | Eub 338f | ACTCCTACGGGAGGCAGCAG | 180 | 1 | 5 | (Bakke et al., 2011) | 95°C-10min; 40 cycles of 95°C-30s, 53°C-40s, 72°C-40s; 95°C-15s, 53°C-1min, 95°C-15s |
| | | Eub 518r | ATTACCGCGGCTGCTGG | | | | | |
| *16S rRNA* Archaea | DSM 23604 *Methanolinea mesophila* | ARC787f | ATTAG ATACC CSBGT AGTCC | 273 | 1 | 5 | (Yu et al., 2005) | 95ºC-10min; 40 cycles of 95ºC-15 s, 57ºC-20s, 72ºC-30s; 95°C-15s, 57°C-1min, 95°C-15s |
| | | ARC1059r | GCCAT GCACC WCCTC T | | | | | |
| *nifH* | DSM 17167 *Paraburkholderia phymatum* | F | AAAGGYGGWATCGGYAARTCCACCAC | 457 | 1 | 5 | (Wallenstein and Vilgalys, 2005) | 95°C-10min; 40 cycles of 95°C-1min, 53°C-27s, 72°C-1min; 95°C-15s, 53°C-1min, 95°C-15s |
| | | R | TTGTTSGCSGCRTACATSGCCATCAT | | | | | |
| *amoA* Bacteria | DSM 28437 *Nitrosomonas europaea* | *amoB* 1F | GGGGTTTCTACTGGTGGT | 491 | 1 | 0.2 | (Rotthauwe et al., 1997) | 95°C-10min; 40 cycles of 95°C-45s, 60°C-45s, 72°C-45s; 95°C-15s, 60°C-1min, 95°C-15s |
| | | *amoB* 2R | CCCCTCKGSAAAGCCTTCTTC | | | | | |
| *amoA* Archaea | *Nitrososphaera viennensis* | *amoA* 1F | STAATGGTCTGGCTTAGACG | 635 | 0.56 | 0.7 | (Francis et al., 2005) | 95°C-5min, 40 cycles of 95°C-40s, 56°C-30s, 72°C-1min; 95°C-15s, 56°C-1min, 95°C-15s |
| | | *amoA* 2R | GCGGCCATCCATCTGTATGT | | | | | |
| *nir*S | DSMZ 1690 *Nitrospirillum brasilense* Sp7 | 4 QF | GTSAACGYSAAGGARACSGG | 410 | 0.67 | 0.3 | (Kandeler et al., 2006) | 95°C-10min; 6 cycles of 95°C-15s, 63°C-30s, 72°C-40s; 38 cycles of 95°C-15s, 58°C–30s, 72°C–40 s; 95°C-15s, 58°C-1min, 95°C-15s |
| | | 6 QR | GASTTCGGRTGSGTCTTSAYGAA | | | | | |
| *nosZ* | BR 11003 *Nitrospirillum brasilensi* | 2F | CGCRACGGCAASAAGGTSMSSGT | 267 | 1 | 5 | (Henry et al., 2006) | 95°C-10 min; 4 cycles of 95°C- 20s, 63°C-30s, 72°C-30s; 40 cycles of 95°C-20s, 60ºC-20s, 72°C–30s; 95ºC-15s, 60ºC-1min, 95ºC-15s |
| | | 2R | CAKRTGCAKSGCRTGGCAGAA | | | | | |

***DSMZ – reference code for strains from the Leibniz Institute DSMZ (Deutsche Sammlung von Mikroorganismen und Zellkulturen GmbH); BR – reference code for strains from the Biological Resources Center Johanna Döbereiner (CRB-JD), Embrapa Agrobiologia, Rio de Janeiro/Brazil, used to construct standard curves for qPCR.**

**Supplementary Table 2 –** Chemical attributes of the organic layers of the forest and pasture. The values represent the mean and standard deviation of five replicates.

| Soil chemical properties | Unity | Forest | | | | | Pasture | |
|---|---|---|---|---|---|---|---|---|
| | | L | F | FHc | $H_1$ | $H_2$ | L | F |
| N | g $kg^{-1}$ | 13.1 ± 2.5 | 16.3 ± 2.6 | 18.8 ± 2.9 | 12.8 ± 5.6 | 12.6 ± 2.6 | 7.35 ± 1.2 | 7.5 ± 2.3 |
| P | g $kg^{-1}$ | 272.2 ± 102.3 | 318.8 ± 75.1 | 294.0 ± 67.1 | 156.7 ± 64.7 | 126.6 ± 16.2 | 192.4 ± 45.3 | 182.6 ± 37 |
| K | mg $kg^{-1}$ | 1.2 ± 0.5 | 1.1 ± 0.4 | 0.8 ± 0.2 | 0.3 ± 0.08 | 0.2 ± 0.01 | 0.4 ± 0.09 | 0.3 ± 0.05 |
| Ca | mg $kg^{-1}$ | 5325 ± 1670 | 4145 ± 1027 | 1576 ± 857 | 301 ± 35 | 273 ± 148 | 5195 ± 504 | 3840 ± 245 |
| Mg | mg $kg^{-1}$ | 2054 ± 688.3 | 1638 ± 716 | 803 ± 375 | 282 ± 182 | 210 ± 148 | 1298 ± 504 | 797 ± 225 |
| Cu | mg $kg^{-1}$ | 85.6 ± 93.9 | 4.9 ± 1.3 | 4.02 ± 1.3 | 1.5 ± 0.5 | 0.26 ± 0.6 | 5.13 ± 2.3 | 3.42 ± 0.9 |
| Fe | mg $kg^{-1}$ | 149.2 ± 45.8 | 269.6 ± 47.4 | 398.80 ± 75.3 | 393.40 ± 156.5 | 383.40 ± 117 | 344.40 ± 81.8 | 2514.4 ± 899.9 |
| Mn | mg $kg^{-1}$ | 90.3 ± 29.5 | 91.9 ± 37.5 | 60.24 ± 21.1 | 17.66 ± 3.9 | 8.91 ± 3.5 | 99.34 ± 26.4 | 94.98 ± 24 |
| Zn | mg $kg^{-1}$ | 16.5 ± 2.5 | 19.6 ± 7.9 | 17.80 ± 6.2 | 8.29 ± 3.3 | 7.21 ± 2.2 | 28.74 ± 3.2 | 21.08 ± 4 |

**Supplementary Table 3 –** Chemical attributes of the organomineral layers of the forest and pasture. The values represent the mean and standard deviation of five replicates.

| Soil chemical properties | Unity | Forest | | Pasture | |
|---|---|---|---|---|---|
| | | $H_3$ | Mineral soil | Mineral soil | Rhizospheric soil |
| C org | g $kg^{-1}$ | 82.9 ± 18.1 | 15.86 ± 3.7 | 10.62 ± 1.3 | 17.98 ± 3.8 |
| N | g $kg^{-1}$ | 5.6 ± 1.0 | 1.16 ± 0.4 | 0.96 ± 0.15 | 1.48 ± 0.3 |
| P | mg $kg^{-1}$ | 7.4 ± 1.5 | 2.40 ± 1.5 | 2.60 ± 2.07 | 4.40 ± 0.9 |
| $Ca^{2+}$ | cmol $kg^{-1}$ | 0.8 ± 0.3 | 0.22 ± 0.08 | 1.24 ± 0.4 | 2.08 ± 0.6 |
| $Mg^{2+}$ | cmol $kg^{-1}$ | 1.0 ± 0.4 | 0.22 ± 0.005 | 0.54 ± 0.4 | 0.8 ± 0.2 |
| $K^{+}$ | cmol $kg^{-1}$ | 0.16 ± 0.05 | 0.02 ± 0.005 | 0.03 ± 0.01 | 0.09 ± 0.02 |
| $Na^{+}$ | cmol $kg^{-1}$ | 0.18 ± 0.11 | 0.04 ± 0.01 | 0.02 ± 0.004 | 0.02 ± 0.005 |
| Cu | mg $kg^{-1}$ | 0.12 ± 0.03 | 0.07 ± 0.08 | 0.11 ± 0.04 | 0.11 ± 0.07 |
| Fe | mg $kg^{-1}$ | 8.13 ±1.3 | 3.96 ± 0.9 | 17.38 ± 4.9 | 22.90 ± 5.1 |
| Mn | mg $kg^{-1}$ | 3.46 ± 2.8 | 0.34 ± 0.3 | 9.58 ± 2.5 | 4.81 ± 1.1 |
| Zn | mg $kg^{-1}$ | 5.30 ± 3.4 | 0.72 ± 0.2 | 2.00 ± 0.7 | 0.93 ± 0.2 |
| $Al^{3+}$ | cmol $kg^{-1}$ | 3.94 ± 0.9 | 1.04 ± 0.4 | 0.01 ± 0 | 0 |
| pH ($H_2O$) | - | 3.82 ± 0.2 | 4.3 ± 0.3 | 5.68 ± 0.4 | 5.76 ± 0.2 |

**Supplementary Figure 1** – Sampling sites and sampling scheme.

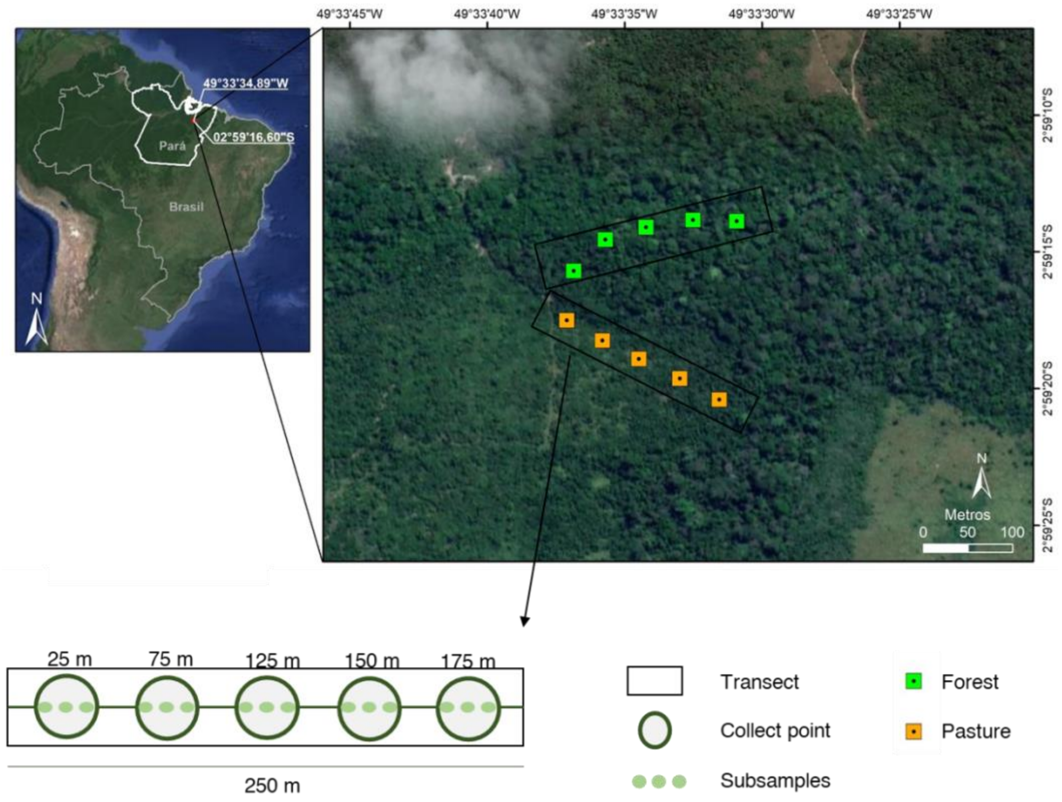


Google Earth (July 2018).

**Supplementary Figure 2.** The C:N ratio, $\delta^{13}C$ and $\delta^{15}N$ of organic material from the layers of forest (L, F, FHc, $H_1c$, $H_1$, $H_2$, $H_3$ and mineral soil) and pasture floors (L, F and mineral soil). RS means the pasture rhizosphere soil. Bars are confidence intervals at 95% probability.

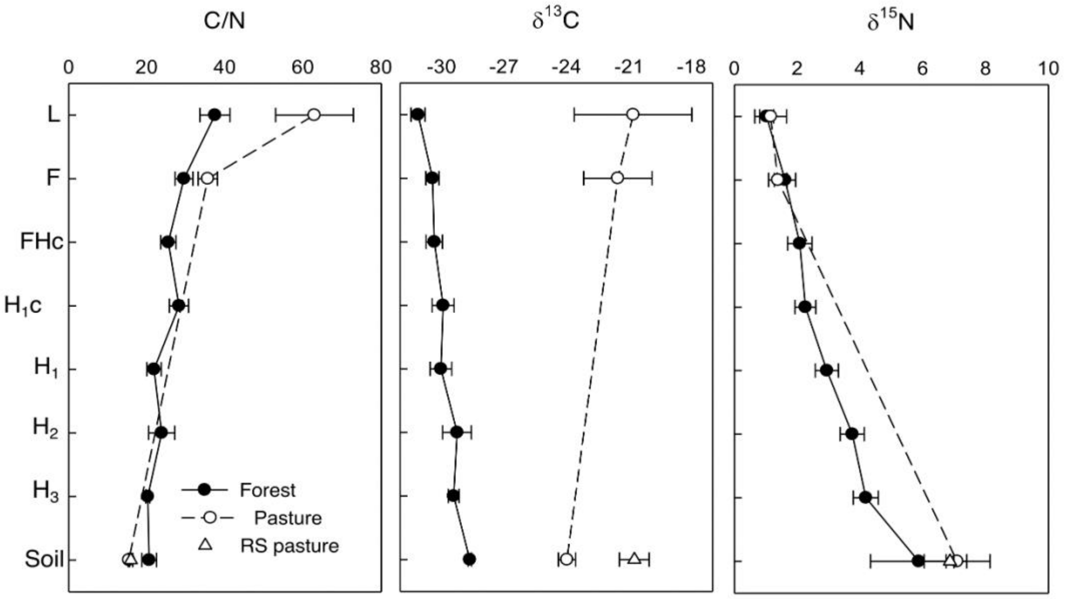

**Supplementary Figure 3** – Detrended correspondence analysis (DCA) of bacterial communities in the organic layers in different stages of decomposition and mineral soil of forest (A, C) and pasture (B, D). Coenoclines in the forest (E) and pasture (F) floors. L, entire leaves; F, fragmented leaves; FH, fine fragments of the mixture of fragmented leaves and humus; FHc, fragments greater than 2 mm from the mixture of fragmented leaves and humus; $H_1$, humus with fine roots; $H_2$, humus; $H_3$, humus mixed with soil mineral material; Rhizo, rhizospheric soil; Soil, initial 10 cm of mineral soil from forest or pasture.

A

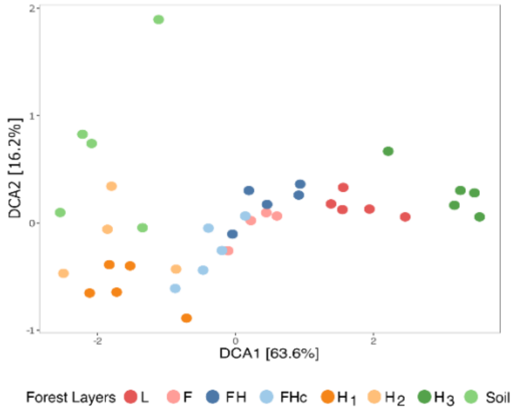


B

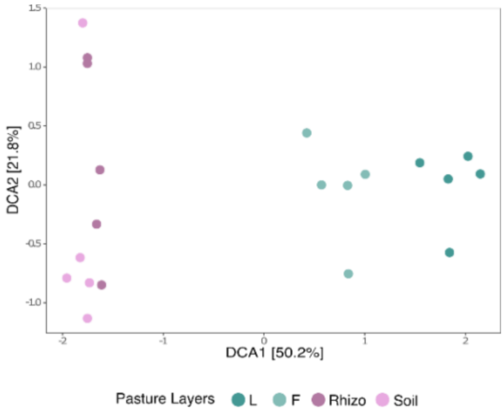


C

| Forest | | | | |
|---|---|---|---|---|
| | DCA1 | DCA2 | DCA3 | DCA4 |
| Eigenvalues | 0.7481 | 0.1903 | 0.1256 | 0.1125 |
| Decorana values | 0.7679 | 0.2089 | 0.1393 | 0.0837 |
| Axis lengths | 6.0828 | 2.7824 | 1.7234 | 1.509 |

D

| Pasture | | | | |
|---|---|---|---|---|
| | DCA1 | DCA2 | DCA3 | DCA4 |
| Eigenvalues | 0.7711 | 0.3348 | 0.2834 | 0.1474 |
| Decorana values | 0.8028 | 0.3715 | 0.2193 | 0.1275 |
| Axis lengths | 4.6768 | 2.5667 | 2.2644 | 1.6525 |

E

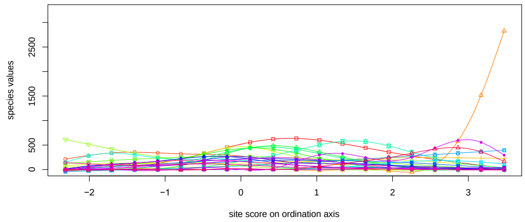


F

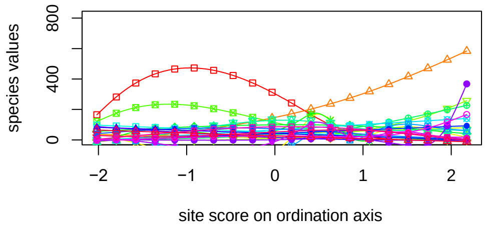

**Supplementary Figure 4** – Relative abundance of the main bacterial phyla in the organic layers and mineral soil of the forest and pasture. Bars indicate the standard deviation. Different letters represent a significant difference at 5% by the Kruskal Wallis test. L, entire leaves; F, fragmented leaves; FH, fine fragments of the mixture of fragmented leaves and humus; FHc, fragments greater than 2 mm from the mixture of fragmented leaves and humus; $H_1$, humus with fine roots; $H_2$, humus; $H_3$, humus mixed with soil mineral material; Rhizo, rhizospheric soil; Soil, initial 10 cm of mineral soil from forest or pasture.

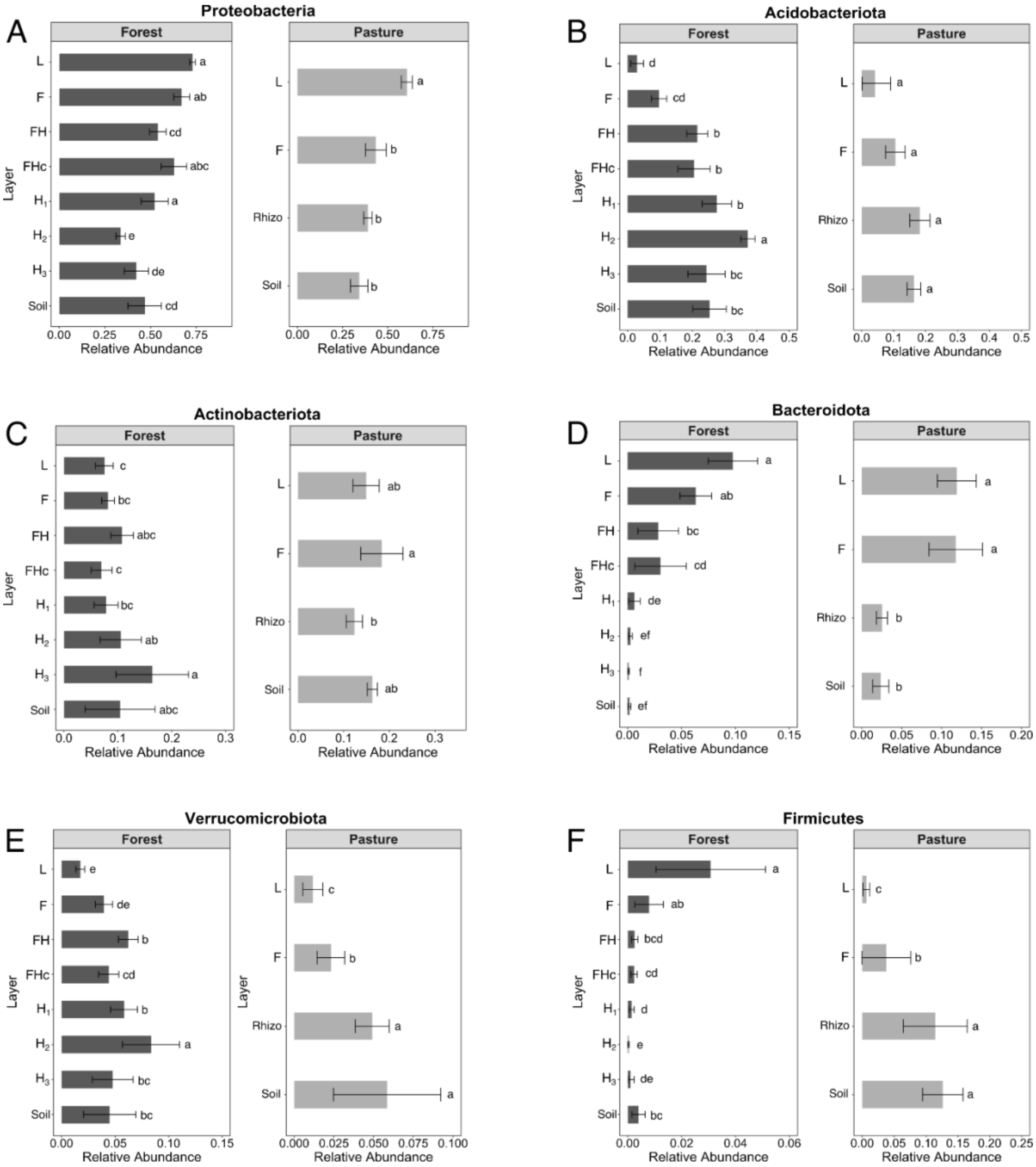

**Supplementary figure 5** – Relative abundance of bacterial classes in organic layers in different stages of decomposition and mineral soil of forest and pasture. L, entire leaves; F, fragmented leaves; FH, fine fragments of the mixture of fragmented leaves and humus; FHc, fragments greater than 2 mm from the mixture of fragmented leaves and humus; $H_1$, humus with fine roots; $H_2$, humus; $H_3$, humus mixed with soil mineral material; Rhizo, rhizospheric soil; Soil, initial 10 cm of mineral soil from forest or pasture.

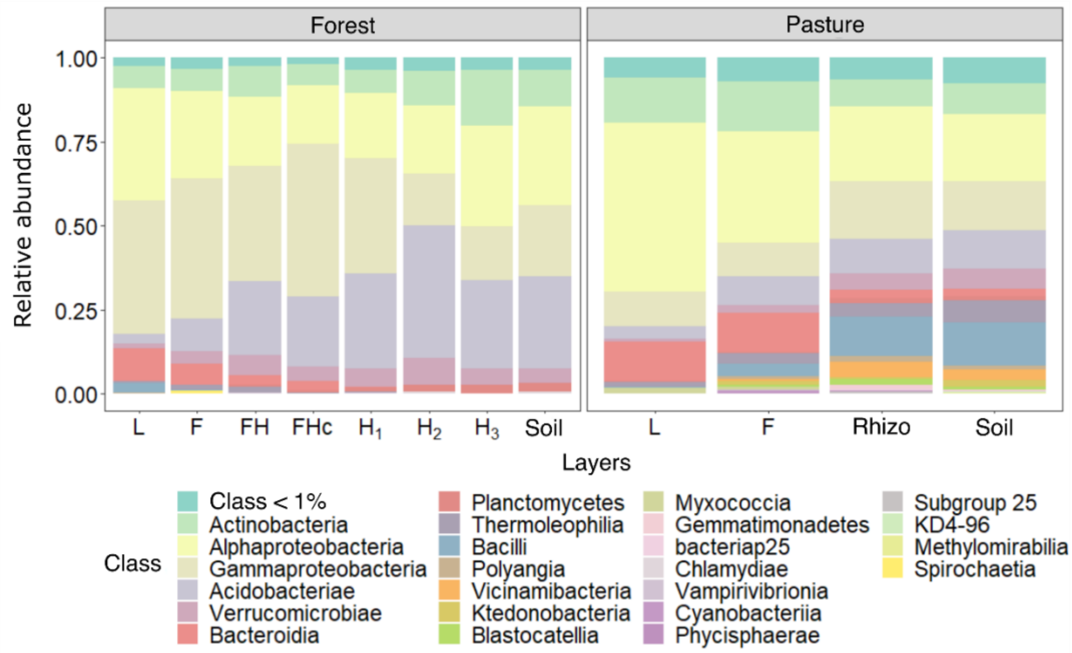

**Supplementary Figure 6** – Relative abundance of bacterial orders in organic layers at different stages of decomposition and mineral soil of forest and pasture. L, entire leaves; F, fragmented leaves; FH, fine fragments of the mixture of fragmented leaves and humus; FHc, fragments greater than 2 mm from the mixture of fragmented leaves and humus; $H_1$, humus with fine roots; $H_2$, humus; $H_3$, humus mixed with soil mineral material; Rhizo, rhizospheric soil; Soil, initial 10 cm of mineral soil from forest or pasture.

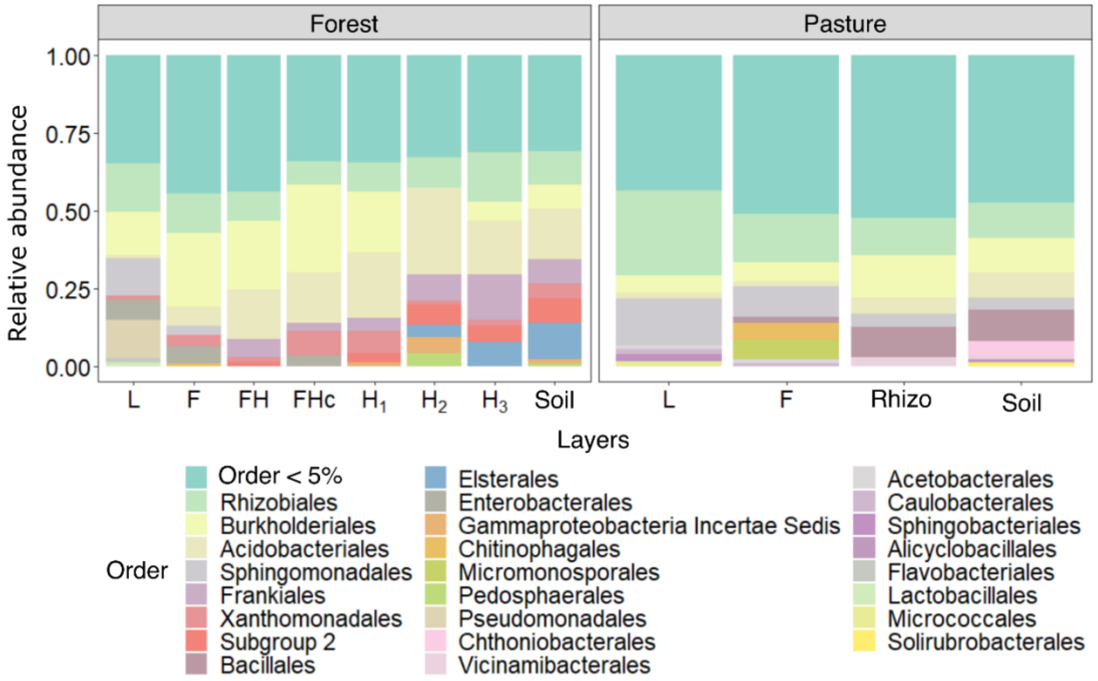

**Supplementary Figure 7** – Relative abundance families in organic layers at different stages of decomposition and mineral soil of forest and pasture. L, entire leaves; F, fragmented leaves; FH, fine fragments of the mixture of fragmented leaves and humus; FHc, fragments greater than 2 mm from the mixture of fragmented leaves and humus; $H_1$, humus with fine roots; $H_2$, humus; $H_3$, humus mixed with soil mineral material; Rhizo, rhizospheric soil; Soil, initial 10 cm of mineral soil from forest or pasture.

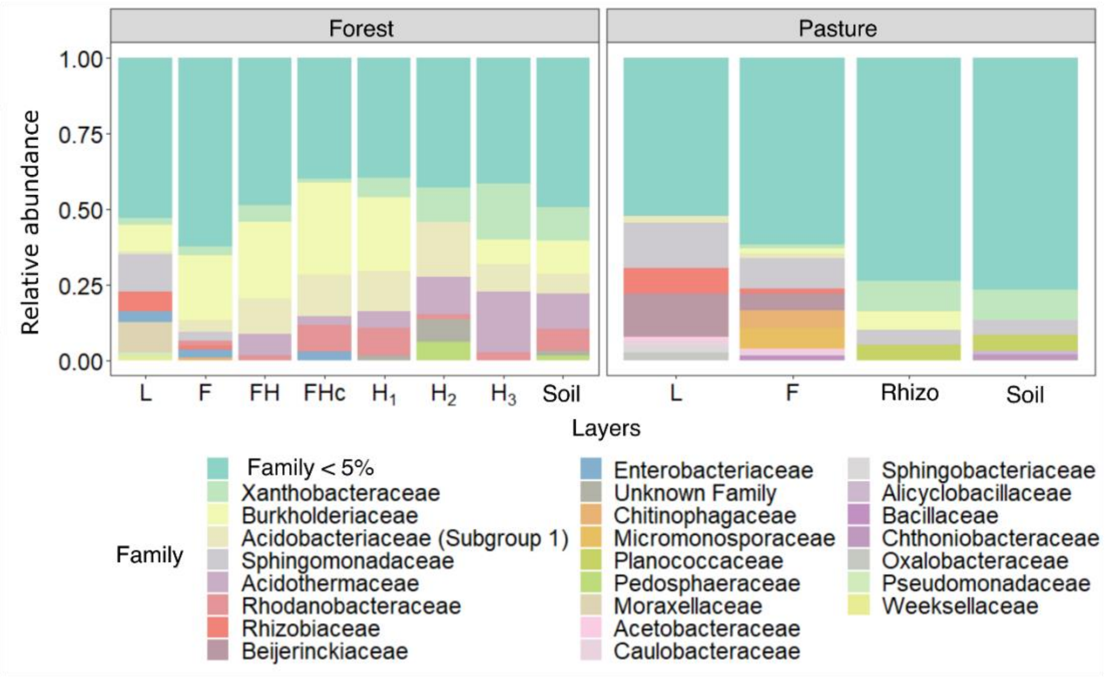

**Supplementary Figure 8** –nirS/nosZ ratios in organic layers in different stages of decomposition and in mineral soil of forest (A) and pasture (B). L, entire leaves; F, fragmented leaves; FH, fine fragments of the mixture of fragmented leaves and humus; FHc, fragments greater than 2 mm from the mixture of fragmented leaves and humus; $H_1$, humus with fine roots; $H_2$, humus; $H_3$, humus mixed with soil mineral material; Rhizo, rhizospheric soil; Soil, initial 10 cm of mineral soil from forest or pasture.

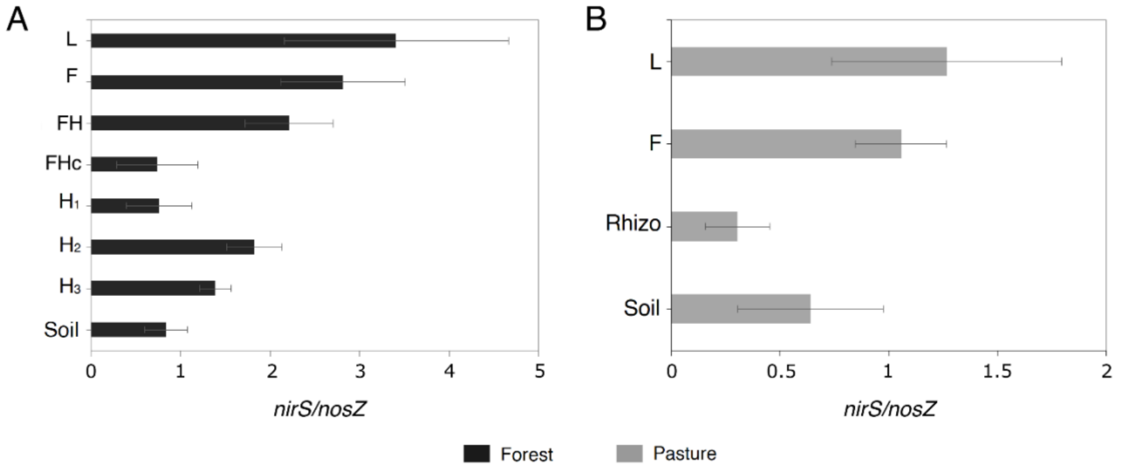


### References

Bakke, I., De Schryver, P., Boon, N., Vadstein, O., 2011. PCR-based community structure studies of Bacteria associated with eukaryotic organisms: A simple PCR strategy to avoid co-amplification of eukaryotic DNA. J. Microbiol. Methods 84, 349–351. https://doi.org/10.1016/J.MIMET.2010.12.015

Empresa Brasileira de Pesquisa Agropecuária – EMBRAPA. 1997. Manual de métodos de análise de solo. 2sd ed. Centro Nacional de Pesquisas de Solos, Rio de Janeiro.

Francis, C.A., Roberts, K.J., Beman, J.M., Santoro, A.E., Oakley, B.B., 2005. Ubiquity and diversity of ammonia-oxidizing archaea in water columns and sediments of the ocean. Proc. Natl. Acad. Sci.U.S.A. 102, 14683–14688. https://doi.org/10.1073/PNAS.0506625102/SUPPL_FILE/06625FIG3.PDF

Henry, S., Bru, D., Stres, B., Hallet, S., Philippot, L., 2006. Quantitative detection of the nosZ gene, encoding nitrous oxide reductase, and comparison of the abundances of 16S rRNA, narG, nirK, and nosZ genes in soils. Appl. Environ. Microbiol. 72,

5181–5189.

Kandeler, E., Deiglmayr, K., Tscherko, D., Bru, D., Philippot, L., 2006. Abundance of narG, nirS, nirK, and nosZ genes of denitrifying bacteria during primary successions of a glacier foreland. Appl. Environ. Microbiol. 72, 5957–5962.

Rotthauwe, J.H., Witzel, K.P., Liesack, W., 1997. The ammonia monooxygenase structural gene amoA as a functional marker: molecular fine-scale analysis of natural ammonia-oxidizing populations. Appl. Environ. Microbiol. 63, 4704–4712. https://doi.org/10.1128/AEM.63.12.4704-4712.1997

Santos, R.D., Jacomine, P.T.K., Anjos, L.H.C., Oliveira, V.A., Lumbreras, J.F., Coelho, M.R., Cunha, T.J.F., Oliveira, J.B., 2018. Sistema Brasileiro de Classificação de Solos, 5th ed. Brasília, DF.

Santos, R.D., Santos, H.G., Ker, J.C., Anjos, L.H.C., Shimzu, S.H., 2015. Manual de descrição e coleta de solo no campo, 7°. ed. Viçisa, MG.

Wallenstein, M.D., Vilgalys, R.J., 2005. Quantitative analyses of nitrogen cycling genes in soils. Pedobiologia (Jena). 49, 665–672. https://doi.org/10.1016/J.PEDOBI.2005.05.005

Yu, Y., Lee, C., Kim, J., Hwang, S., 2005. Group-specific primer and probe sets to detect methanogenic communities using quantitative real-time polymerase chain reaction. Biotechnol. Bioeng. 89, 670–679. https://doi.org/10.1002/BIT.20347
